# Untangling the contribution of adaptive *versus* non-adaptive processes in the evolution of reproductive isolation between *Coenonympha* butterflies

**DOI:** 10.1101/2024.11.29.625973

**Authors:** Thibaut Capblancq, Camille Roux, Frédéric Boyer, Fabrice Legeai, Mathieu Joron, Laurence Després

## Abstract

Speciation is a key evolutionary process which has been studied in numerous organisms and at multiple scales, from lineage radiation to gene expression. However, the factors explaining the rise of new species are not yet fully understood, and the relative contribution of neutral *versus* selective evolutionary processes in triggering and maintaining reproductive isolation between lineages is still debated. To explore this question, we study the divergence of two butterfly species, *Coenonympha arcania* and *C. gardetta* (Nymphalidae), which diverged relatively recently but show strong ecological differences. Whole genome sequence data reveal high overall differentiation between the two lineages, best explained by a long period of isolation at the early stage of their divergence. Demographically explicit approaches identify that 6.6% of the genome (32.7 Mbp) is impermeable to gene flow between the two species. Lots of these barrier loci are located on the Z chromosome, potentially spanning 75% of its length which would indicate that a large Z effect is at play in this speciation. Moreover, only a small proportion of barriers showed signatures of selection, suggesting that non-adaptive processes largely contributed to the build-up of reproductive isolation. Therefore, although genes involved in stress response and response to hypoxia are interesting candidates under selection, the adaptation of *C. gardetta* to alpine conditions may not to be the main driver of speciation. Our study brings an original example of intertwined adaptive and non-adaptive processes leading to reproductive isolation in a speciation with secondary contact and improves our understanding of the genomic underpinnings of species divergence.

## INTRODUCTION

The extensive array of living organisms emerges from a fundamental process in evolution: speciation. In the 20th century, this process was regarded as a series of binary divisions in evolutionary lineages, primarily depicted through phylogenetic trees. While this dichotomous perspective upholds the evolutionary connections between groups, its significance diminishes when lineages are closely related. Hence, speciation is now well accepted as a continuous process, gradually giving rise to distinct entities and eventually species (Stankowski & Ravinet, 2021). The accumulation of mutations in the genomes progressively results in the isolation of the two lineages by reducing the fitness of their hybrids (Presgraves, 2010). Within the whole range of substitutions between two species, only some affect hybrid fitness, either by generating incompatibilities in gene interactions (*i.e.*, endogenous postzygotic isolation), by being involved in ecological adaptations (*i.e.*, exogeneous postzygotic isolation) or in mating signals (*i.e.*, prezygotic isolation) (Coyne & Orr, 2004). The relative importance of these different types of hybrid incompatibility on the evolution of reproductive isolation varies depending on the demographic and ecological contexts of the speciation event.

Identifying the evolutionary processes that led to the establishment of such barriers between closely related lineages is critical for understanding the emergence of new species (Seehausen *et al*., 2014). Their nature can be broadly classified according to whether they originated through adaptive or non-adaptive processes. Non-adaptive forces lead to the formation of barriers when interacting genes co-evolve differently within isolated lineages. Negative interactions between these co-evolving loci have been shown to be commonly expressed in hybrids, leading to strong reproductive barriers (Bateson, 1909; Dobzhansky, 1936; Muller, 1942). In such case, the formation of hybrids leads to the expression of new interactions that have not been proven by natural selection (Simon *et al*., 2018). Each set of co-evolving deleterious and compensatory mutations can thus become a potential endogenous barrier, impacting the hybrid fitness regardless of the ecological context in which the parent species evolved. Conversely, adaptive processes involve loci whose genotype fitness is directly dependent on the environment. In that case, hybrids tend to be less fit than one of the two parents, depending on the environment in which they are located (Schluter, 2009). The relative importance of the above mentioned non-adaptive and adaptive processes in the build-up of reproductive isolation is tightly linked to the demographic and ecological contexts in which the speciation takes place. In sympatry with environmental heterogeneity, the predominant drivers of barriers are adaptive processes and exogeneous selection (Nosil, 2012), while in allopatry endogenous barriers may arise as a result of genetic drift (Endler, 1977; Barton & Hewitt, 1985; Noor & Bennett, 2009).

To investigate empirically the comparative influences of adaptive and non-adaptive processes on reproductive isolation, it is essential first to pinpoint the genomic regions linked to these barriers, the so-called “barrier loci” (Roux *et al*., 2016; Ravinet *et al*., 2017). This step allows analyzing the evolutionary forces that have influenced their recent history. Identifying species barriers *in natura* can be achieved when a pair of closely related species are experiencing gene flow, either as a result of secondary contact, or continuously during their divergence process (Smadja & Butlin, 2011; Martin *et al*., 2013). If strong selection acts against migrant alleles in specific genomic barriers while the rest of the genome is exchanged more freely between species, then patterns of nucleotide polymorphism and divergence may be sufficiently contrasted to detect barrier loci (Fraïsse *et al*., 2021; Laetsch *et al*., 2023). In that spirit, new methodological developments have taken advantage of large genomic and computational resources to test a local reduction in migration rate using window-based approaches. Two methods have recently been distributed to carry out genomic scans of species barriers: gIMble (Laetsch *et al*., 2023) and DILS (Fraïsse *et al*., 2021). These methods have the advantage to account for the confounding effect of background selection, which can produce differentiation patterns similar to those of divergent selection (Charlesworth *et al*., 1997; Cruickshank & Hahn, 2014). Background selection results in a local reduction of genetic diversity due to purifying selection against deleterious mutations, and promotes high species differentiation without playing a functional role in reproductive isolation (Charlesworth, 2012). The two above mentioned approaches tackle this issue and show great potential to identify genomic regions involved in reproductive isolation. The identification of such barriers stands as a prerequisite to unravel the evolutionary mechanisms determining their evolution. For instance, contrasting the genomic patterns of polymorphism and divergence within these barrier regions against the broader genomic background allows testing whether they are predominantly shaped by local adaptive processes or not (McDonald & Kreitman, 1991; Stoletzki & Eyre-Walker, 2011; Han *et al*., 2017; Irwin *et al*., 2018; Shang *et al*., 2021; Moreira *et al*., 2023).

In this study we dissect the genetic architecture of reproductive isolation between two temperate butterflies, *Coenonympha arcania* and *C. gardetta*, and estimate the relative contributions of adaptive *versus* non-adaptive processes to their speciation. The two species are ecologically divergent, with *C. arcania* being widely distributed in Europe, from sea level to elevations around 1,500 meters, while *C. gardetta* is typically encountered in subalpine and alpine habitats, above 1,500 meters in the Alps. They diverged around 1.7 million years ago (Capblancq *et al*., 2015), are morphologically distinct and have parapatric distributions with multiple contact zones at intermediate elevation around the Alps. The two species do not form two distinct monophyletic groups based on mtDNA, which suggests that mitochondrial transfers (and gene flow) occurred sometime during their divergence (Capblancq, 2016). Occasional hybrids between the two species can be observed in regions of sympatry (Capblancq *et al*., 2019), confirming a potential for ongoing gene flow, although they would supposedly be unfit or mostly sterile according to previous experimental crosses (de Lesse, 1960). This strong but incomplete reproductive isolation makes the species pair an ideal model to study the genomics of speciation and identify barrier loci along the species genome. In presence of gene flow, if ecological divergence is the main driver of speciation (i.e., ecological speciation), loci involved in local adaptation and reproductive isolation should be largely shared. Conversely, if reproductive isolation evolved independently from local adaptation, signatures of restricted gene flow should not be necessarily associated with signature of selection.

To explore these questions, we surveyed whole genomes from natural populations of the two species with four main objectives: 1) study the temporal patterns of gene flow between *C. arcania* and *C. gardetta* (*i.e.*, isolation *versus* continuous migration *versus* secondary contact *versus* ancestral migration), 2) locate barrier loci where gene flow is effectively blocked, 3) evaluate the relative importance of adaptive *versus* non-adaptive processes in determining reproductive isolation, and 4) identify candidate speciation genes. To achieve this, we generated whole-genome short read data for 19 individuals of *C. arcania* and 17 individuals of *C. gardetta* and mapped the sequences to the *C. arcania* reference genome (Legeai *et al*., 2024). We then explored genetic diversity and differentiation along the genome of the species using 50kbp windows and located genomic regions displaying reduced gene flow (referred to as “barrier loci”) using both gIMble and DILS. We then sought signatures of selection both within and outside these barrier regions, in order to qualify the forces shaping the observed variation in gene flow levels along the genome.

## RESULTS

A total of 65.2 million cleaned reads were obtained on average per individual (range: 46.6-88.5 million reads), of which 54.5% mapped to the *C. arcania* reference with a quality score equals or above 20. After removal of PCR duplicates, mean coverage across all individuals was 18x (range: 13-25.2x), and genotype likelihoods were obtained for 203,075,624 sites that passed quality and coverage thresholds for both species, scattered across 9,844 50kb genomic windows representing 99% of the reference genome length (497Mbp, Legeai *et al*., 2024). Among these sites, 16,571,294 were polymorphic (8.2% of the covered sites) and were homogeneously distributed along the 30 first large scaffolds of the reference (Supplemental Fig. S1).

### Demographic inferences favor a scenario of secondary contact and low levels of gene flow between species

To investigate the history of divergence and gene flow between *C. arcania* and *C. gardetta* we compared the likelihood of four speciation scenarios: strict isolation (SI), isolation with migration (IM), secondary contact (SC) and ancient migration (AM), using Fastsimcoal2 (Excoffier *et al*., 2013) and DILS (Fraïsse *et al*., 2021). We used the whole dataset for demographic inferences because including or not the sexual chromosome Z gave very similar results (Supplemental Table S1).

Both procedures favored a model of isolation followed by a secondary contact for the divergence of *C. arcania* and *C. gardetta* (Figure 1), in contrast with previous results that favored an isolation with migration scenario (Capblancq *et al*., 2019), but confirming the presence of gene flow during the speciation process. The initial divergence time estimates were very similar for the two methods with Fastsimcoal2 inferring 1.79 Mya and DILS inferring 1.93 Mya (Figure 1). The secondary contact would have occurred only after a relatively long period of isolation, 360,240 generations ago according to Fastsimcoal2 and 498,562 generations ago according to DILS. The migration rate estimated after the secondary contact between the two species was low (Fastsimcoal2: *m_e_* _arc-gar_ = 9.07e-8, *m_e_* _gar-arc_ = 3.99e-7; DILS: *m_e_* _arc-gar_ = 2.9e-7, *m_e_* _gar-arc_ = 7.9e-7), roughly corresponding to an exchange of 0.5 to 1.5 individuals every generation. Finally, the current *N*_e_ estimates inferred were 2,955,432 (Fastsimcoal2) or 2,659,735 (DILS) for *C. arcania* and 1,748,050 (Fastsimcoal2) or 1,257,299 (DILS) for *C. gardetta*, this time very much consistent with previous estimates inferred from ddRAD-seq data (Capblancq *et al*., 2019).

**Figure 1:**
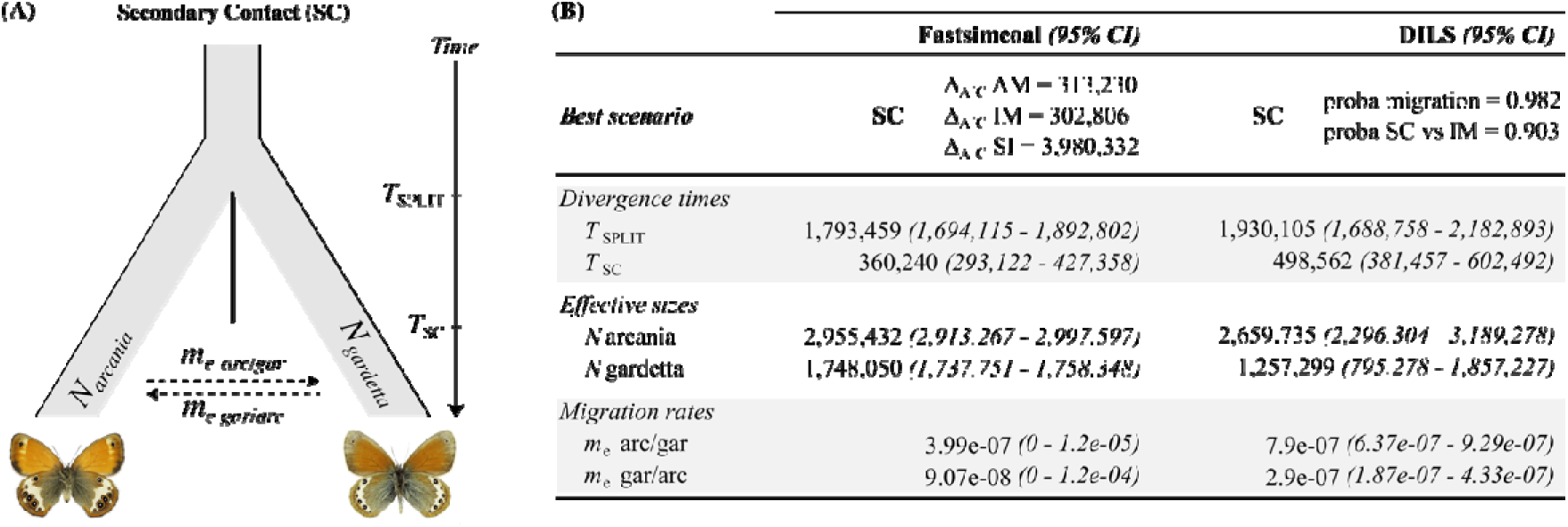
Scenario of divergence between *C. arcania* and *C. gardetta.* (A) Most likely scenario of divergence with its associated parameters. (B) results of Fastsimcoal2 and DILS inferences with the most likely model (SC) in comparison with the three other tested scenarios of divergence (AM, IM and SI) and the inferred parameter values. 95% confidence intervals are shown in brackets.

### Identifying genomic barriers to gene flow unravels the importance of the Z chromosome and several autosomal regions

To locate putative barrier loci, we used two recently published model-based approaches that test for a local reduction of migration along the chromosome of two diverging species: an adaptation of gIMble (Laetsch *et al*., 2023) – the Δ*_B_* method – and DILS (Fraïsse *et al*., 2021). Both approaches follow a very similar process that does not contrast different genomic regions among each other, but two demographic models (with or without gene flow) within each genomic window. The gene flow *vs* no gene flow comparison is made at the window level specifically to account for the variation in local effective population sizes along the genome, variation that can originate from evolutionary factors (linked selection) or be due to intrinsic parameters (reduced recombination rates). The two methods identified 654 genomic regions acting as barrier to gene flow between the genomes of *C. arcania* and *gardetta*, representing 6.6% of the genome. These regions were mostly located on the sex chromosome Z (363 windows = 56% of the barriers) but also on multiple autosomal regions of high genetic differentiation (Figure 2A). The Δ*_B_* method returned 6.5% of the genome as barrier to gene flow, with almost the entire Z chromosome (74.7%), suggesting that the very high *F*_ST_ and low diversity estimates along this sex chromosome (Supplemental Fig. S2) were not only due to its reduced effective size (3/4 of autosomal *N*_e_). We also tried a migration rate threshold inferred only with the Z chromosome to estimate the Δ*_B._* The procedure still returned multiple barrier loci in the Z chromosome but only representing 8.6% of its length (Supplemental Fig. S3). False Positive Rate (FPR) estimated at the window level using simulations was zero for most genomic regions, and only 40 windows returned a non-zero FPR of 1%, supporting the robustness of the procedure. DILS, which, in general, identified a reduced number of barriers compared to the Δ*_B_* method (only 0.3% of the genome), also found nearly half of the barriers (11/27 windows) on the Z chromosome (Figure 2A). The substantial difference between the number of regions identified as barrier by the Δ*_B_* method (642 windows) and DILS (27 windows) was somewhat surprising, with only 15 common windows. This calls for further comparisons of the procedures in the future. Hereafter, we considered as barriers all 654 windows that were identified by either one or the other method, or both. Generally, these windows were localized in genomic regions of increased genetic differentiation and low diversity, but not always (*e.g.,* on chromosome 2, 10 or 12), confirming that genetic differentiation and resistance to gene flow may not always be associated.

**Figure 2:**
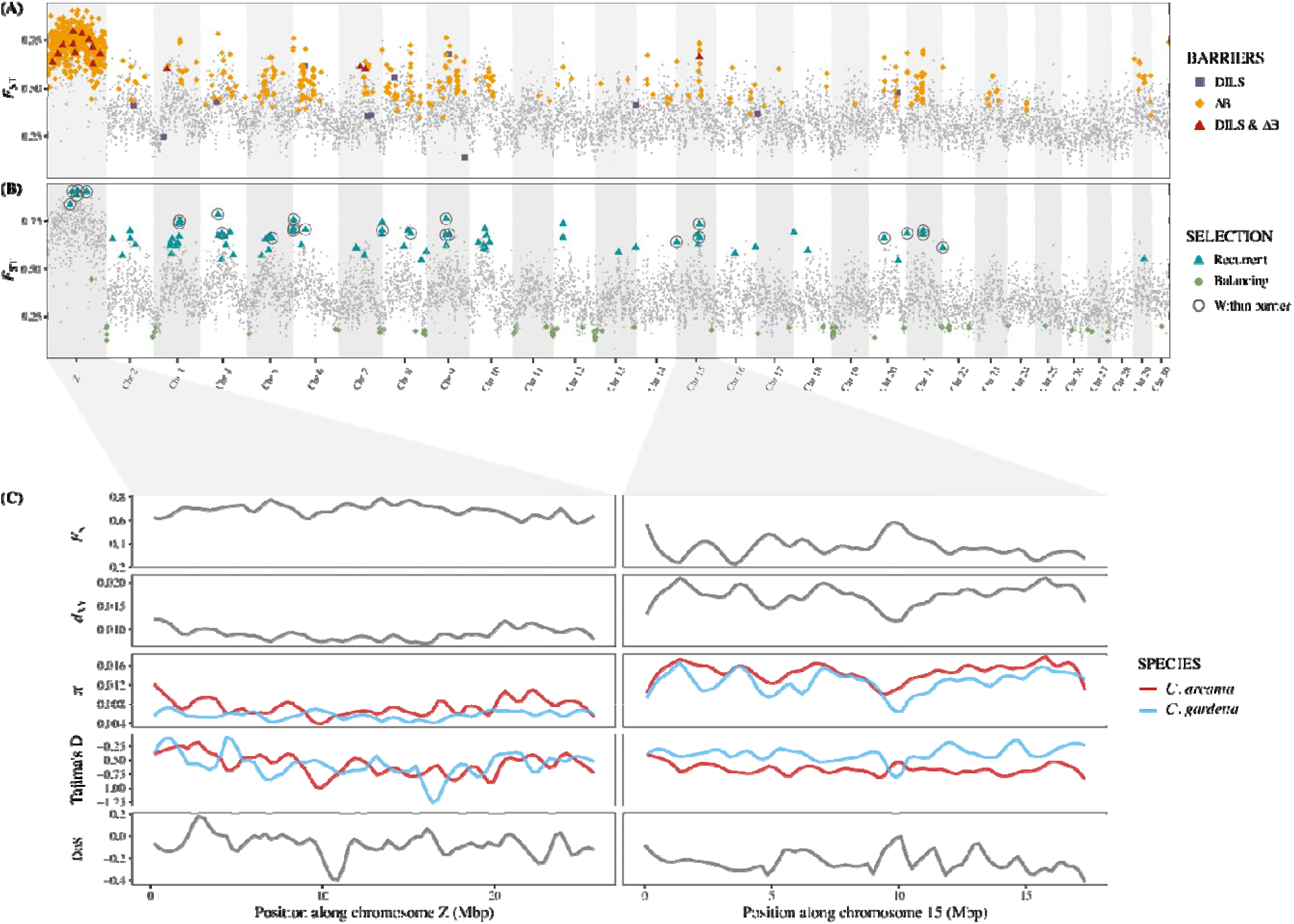
Identification of barriers to gene flow between the genomes of *C. arcania* and *C. gardetta*. (A) Genomic windows considered as resistant to gene flow using two process-based procedures: the Δ*B* procedure (Δ*B* > 0) and/or DILS (posterior probability of being a barrier > 0.7). (B) Windows falling into one of the four selection scenarios described in (Shang *et al*., 2021) based on extreme patterns of *F*_ST_, *d*_XY_ and π. (C) Zoom on chromosome Z and 15 showing different genetic parameters for each genomic window and species. Barriers detected with both DILS and Δ*_B_* approaches are localized with shaded grey rectangles.

Patterns of differentiation (*F*_ST_), divergence (*d*_XY_), and diversity (π) along the two species genomes were also investigated to locate “genomic islands” in the genomic landscape. The procedure of Shang et al (2021) was used, which allows to distinguish three scenarios of selection (recurrent, divergent, balancing) from a neutral signal by identifying extreme combinations of π, *d*_XY_ and *F*_ST_. Only genomic regions putatively under recurrent selection or balancing selection were identified (Figure 2B). The recurrent selection loci, which showed extremely high values of *F*_ST_, low values of π and average values of *d*_XY_, were found in multiple autosomal regions and the Z chromosome. Recurrent selection corresponds to a selective pressure that is acting on the two diverging lineages and was already operating in the ancestral population. The 24 genomic regions located within barriers and identified as under recurrent selection (circled points in Figure 3B) constituted the genomic window “linked to barrier and with signals of positive selection” set for downstream analyses. One of these regions was located on the Z chromosome and the rest distributed on nine of the 29 autosomes.

**Figure 3:**
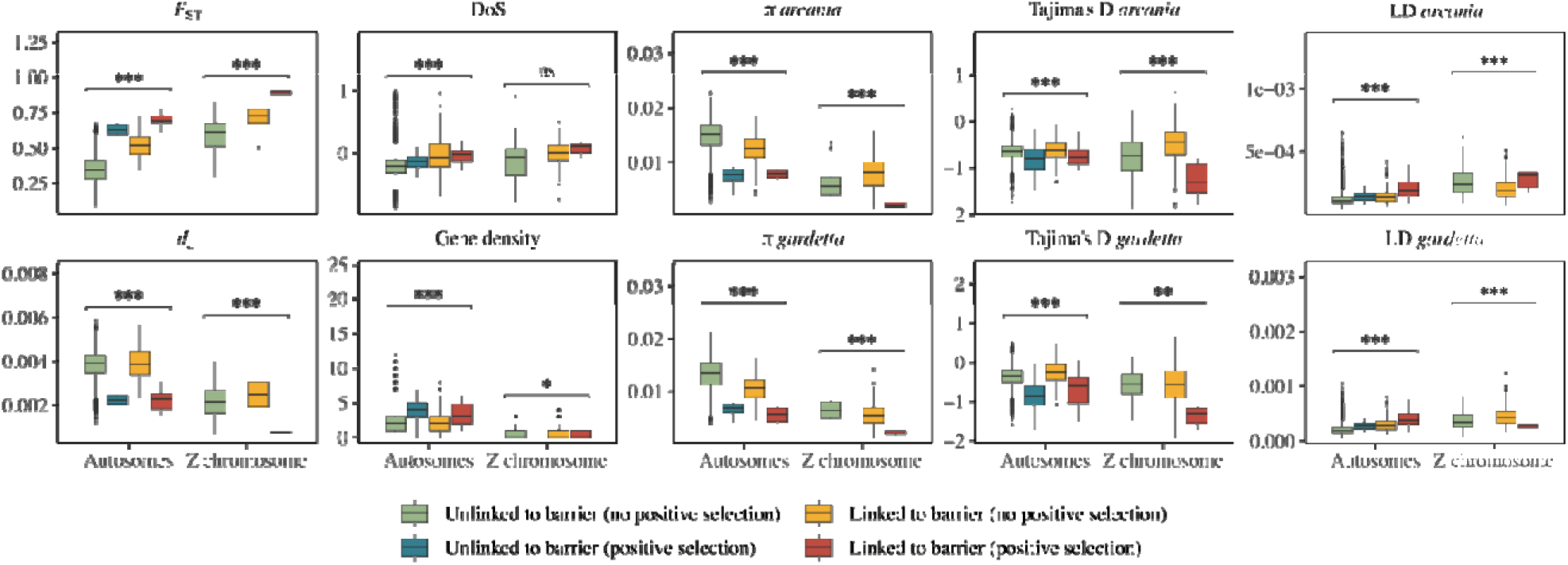
Distribution of genomic parameters in autosomes and chromosome Z, within or outside barrier loci and within or outside windows under positive selection. Are considered barrier loci here all the genomic windows that returned a Δ*_B_* > 0 and/or a DILS posterior probability > 0.7 and fell in one of the genomic islands identified in Figure 2(A). Student’s t-tests returned extremely significant results (*p-value* < 0.001) between the Z chromosome and the autosomes for all parameters. One-way ANOVA tests with the four categories as grouping factor were conducted independently for the Z chromosome and the autosomes for each genomic parameter: a *p-value* < 0.001 is indicated by ‘***’, *p-value* < 0.01 by ‘**’, *p-value* < 0.05 by ‘*’ and ‘ns’ for a *p-value* > 0.05.

Zooming in on these candidate regions often allowed to identify peaks or “islands” in the genomic landscape (Figure 2C). As expected, the candidate regions showed elevated *F*_ST_ and decreased genetic diversity, but they also returned elevated values of DoS (Direction of Selection) in coding regions, suggesting more positive selection than in the rest of the genome. For example, a peak of high DoS values (>> 0) was observed in the middle of chromosome 15 (Figure 2C), a genomic region identified as a barrier to gene flow and exhibiting increased genetic differentiation between and depleted diversity within the two species. However, when DoS is examined gene by gene along the genome, identifying a specific pattern was difficult and many genes with very high DoS values were found in genomic regions not identified as potential barriers or not showing any specific genetic diversity or differentiation (Supplemental Fig. S4).

### Can we disentangle the evolutionary processes shaping heterogenous genomic resistance to gene flow?

To better understand the evolutionary forces that shaped the landscape of genetic diversity within and differentiation between *C. arcania* and *C. gardetta*, multiple genetic parameters were inspected across 9,844 genomic windows on the Z chromosome and the autosomes. As expected, the Z chromosome showed different patterns compared to autosomes (Figure 3, Supplemental Fig. S2), with an *F*_ST_ almost twice as high on Z than on autosomes (*F*_ST(Z)_ = 0.687; *F*_ST(A)=_ 0.357), a much lower net divergence (*d*_a(Z)_ = 0.0024; *d*_a(A)_ = 0.0038), nucleotide diversity twice as low in both species (π_(Z)arcania_= 0.0074; π_(A)arcania_= 0.0145; π_(Z)gardetta_= 0.0058; π_(A)gardetta_ = 0.0131), a much higher DoS on average (DoS_(Z)_ = −0.033; DoS_(A)_ = −-0.201) and a lower density of genes. Tajima’s *D* values were similar on the sex chromosome (Tajima’s *D*_(Z)arcania_ = −0.58; Tajima’s *D*_(Z)gardetta_ = −0.58) but the alpine specialist showed larger estimates on the autosomes (Tajima’s *D*_(Z)arcania_ = −0.65; Tajima’s *D*_(Z)gardetta_ = −0.36). In a similar trend, the genomic windows identified as barriers showed higher differentiation (*F*_ST(barrier)_ = 0.63; *F*_ST(no barrier)_ = 0.37), lower net divergence (*d*_a(barrier)_ = 0.0031; *d*_a(no barrier)_ = 0.0038), higher signal of positive selection (DoS_(barrier)_ = −0.017; DoS_(no barrier)_ = −0.196) and lower genetic diversities (π_(barrier)arcania_ = 0.0097; π_(no barrier)arcania_ = 0.0143; π_(barrier)gardetta_ = 0.0076; π_(no barrier)gardetta_ = 0.0128).

Four categories of genomic regions were distinguished. The genomic windows identified as barriers were distinguished from the non-barrier windows and, within each of these two categories, windows showing signals of positive selection (*i.e.*, recurrent selection in Figure 2) were distinguished from the windows showing no signals of selection. The four resulting types of windows were: i) unlinked to barrier and without signal of positive selection, ii) unlinked to barrier but with signals of positive selection, iii) linked to barrier without positive selection and iv) linked to barrier with positive selection. Among the windows showing no signal of positive selection, net divergence (*d*_a_), gene density and Tajima’s D were similar within and outside barriers, whereas a higher genetic differentiation (*F*_ST_), larger DoS and lower autosomal nucleotide diversity (π) were found in windows linked to barriers compared to those outside barriers (Figure 3). Windows with signals of positive selection, linked or not to barriers, showed increased *F*_ST_, lower *d*_a_, π and Tajima’s D compared to regions with no positive selection, but also a higher density of genes. Significantly higher values of *F*_ST_ (t-test: *p-value* = 3.9e-06) and DoS (t-test: *p-value* = 0.014) were found in windows linked to barriers compared to non-barrier regions when positive selection was identified (Figure 3). However, genomic windows linked to barriers but without selection were not significantly different than regions unlinked to barriers in terms of *d*_a_, DoS or gene density, confirming that their resistance to interspecific gene flow was not linked to undetected positive selection. In the same vein, linkage disequilibrium (LD) was much higher in barriers with positive selection than in the other three categories, consistent with hitchhiking effects. On the contrary, barrier regions with no signal of positive selection returned LD values comparable to non-barrier windows, suggesting little linked selection.

Finally, to avoid a loss of information associated with averaging gene-level information across 50kb genomic window we complemented our analysis with a gene centered approach, by looking at the proportion of genes showing a positive DoS in the four previously described categories (Table 1). Overall, 8.4% of the 15,239 analyzed genes returned a DoS > 0. Windows unlinked to barriers and showing no positive selection were depleted in positive DoS, suggesting a major role of purifying selection in these regions. On the contrary, windows unlinked to barriers but showing signals of selection showed higher values of DoS, presumably because directional selection counteracted purifying selection in these regions. Genomic regions associated with barriers returned an overall inflated proportion of genes with positive DoS and hosted very few genes with extremely low DoS value (< –0.5). This confirmed that the patterns observed for DoS on Figure 3 were not due to a few high DoS outlier genes in windows linked to barriers, but to a more general trend in these regions.

**Table 1:**
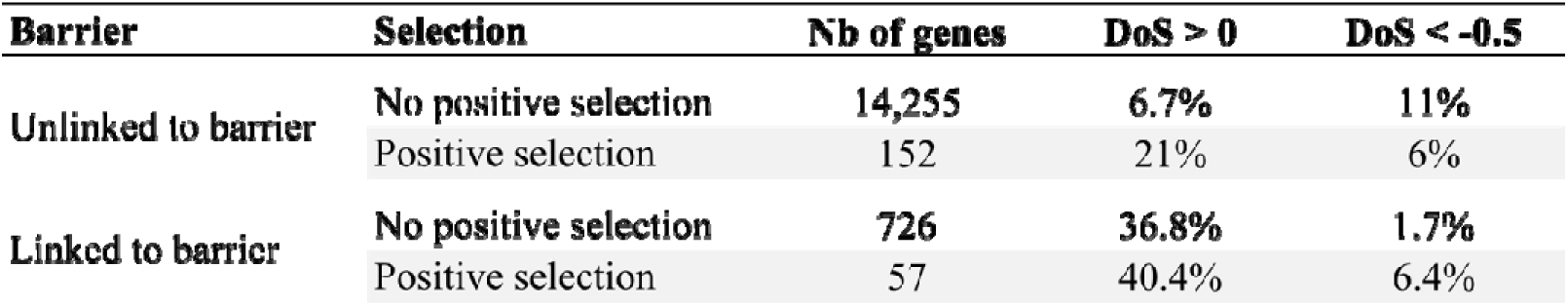
Proportion of genes returning positive (>0) or a very negative (<-0.5) value of Direction of Selection (DoS) within the four different types of genomic windows identified in the study.

### The genes of reproductive isolation (?)

To explore the potential differences in gene functions within the two types of barriers (with or without signal of positive selection), a gene ontology (GO) terms enrichment analysis was conducted for each group. Significantly enriched GO terms were largely shared between the barriers showing signals of positive selection and those showing no such signals (Figure 4). These shared terms included one term associated with energy production: “electron transport chain”, two linked to development: “cell morphogenesis” and “axon guidance”, and another two associated with lipid production: “phospholipid metabolic process” and “lipid biosynthetic process”. One biological process enriched only in barriers with no signals of selection was linked to “sensory perception of smell”, suggesting incompatibilities in the olfactive systems of the two species, without selection acting within or between species. On the contrary, one biological process significantly enriched only in barriers showing signals of positive selection was associated with methylation. The Z chromosome was initially considered separately from the autosomes in the GO terms enrichment analysis, but only one biological process was slightly significantly enriched in the barrier windows of the sex chromosome: “regulation of catalytic activity”; this means that all the processes showing enrichment in barrier windows (Figure 4) are mostly driven by enrichment in autosomal barriers.

**Figure 4:**
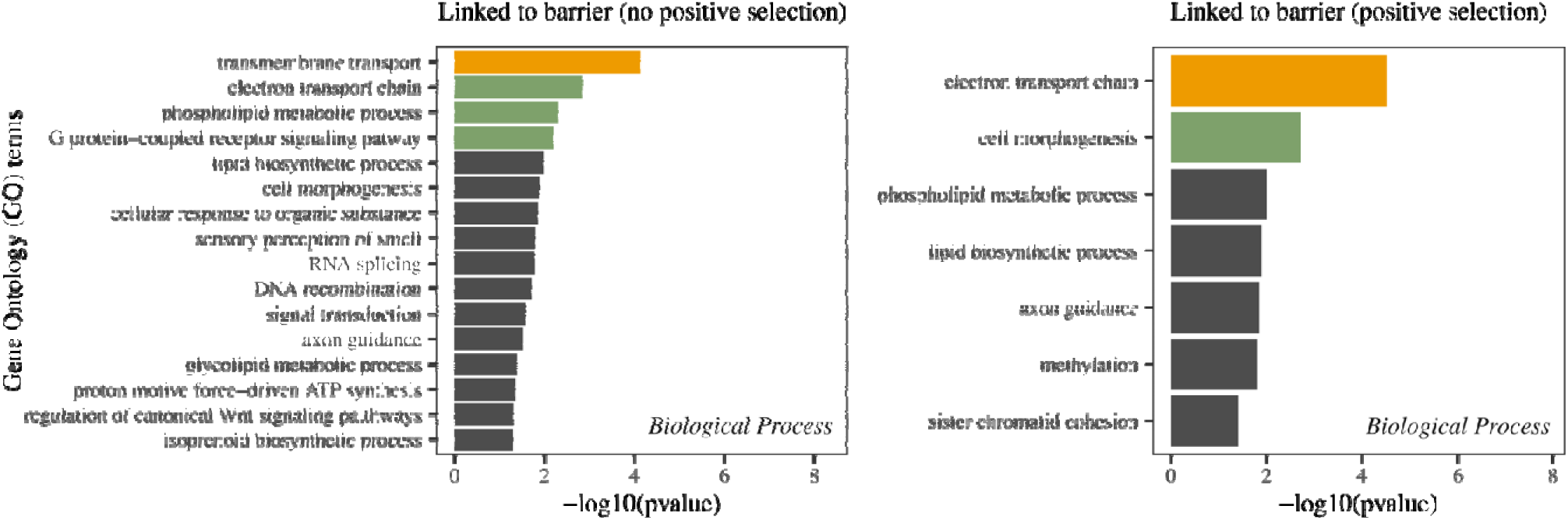
Results of the GO terms enrichment in the genomic regions identified as putative barrier loci between *C. arcania* and *C. gardetta* (both with and without signals of positive selection), with the associated biological processes. Only GO terms found significantly enriched with a Kolmogorov Smirnov test are shown (*p-value* < 0.05) and GO terms shown in green and yellow have strongly significant test results, *p-value* < 0.01 and < 0.001, respectively.

We then inspected the genes located within genomic barriers that also showed signals of positive selection, hypothesizing that they were the ones driving reproductive isolation between *C. arcania* and *gardetta*. In these windows, we found many genes linked to response to stress and especially hypoxia in other species, including those encoding serine/threonine protein kinases, apoptosis-inducing factors, phosphatidylserine synthases or fatty acid synthetases (Table 2).

**Table 2:**
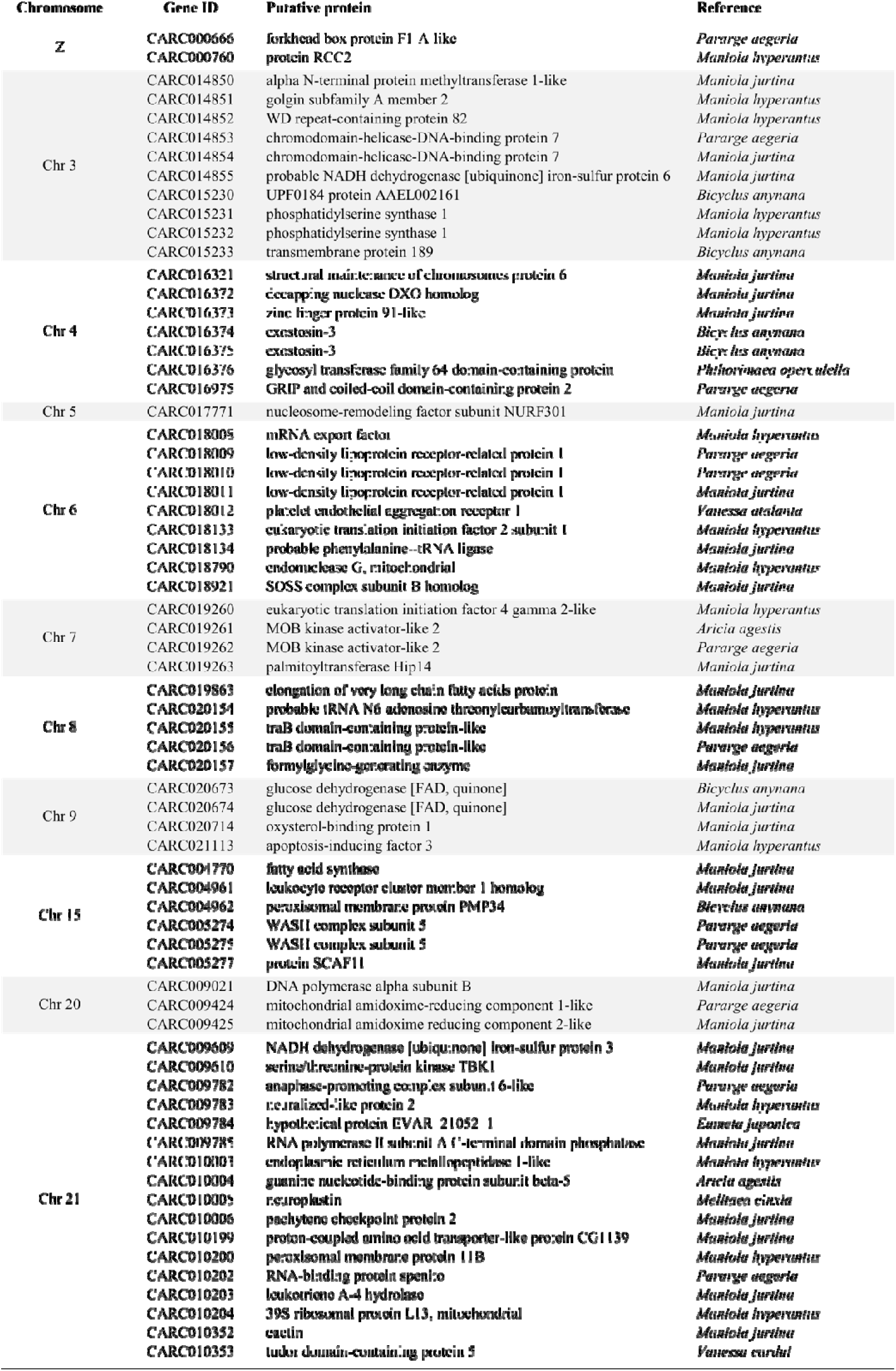
List of the 68 protein-coding genes annotated within the genomic windows identified as barrier loci between *C. arcania* and *C. gardetta* and showing signals of positive selection. The table includes gene IDs, a description of the associated protein functions and the reference organism for which the highest homology was found by Diamond v2.0.13 on NCBI NR 2022-11-30.

## DISCUSSION

When studying divergence between closely related species showing strong ecological divergence in close geographic proximity (*i.e.*, sympatry or parapatry), it is tempting to hypothesize that speciation is mainly driven by divergent selection (Schluter, 2009). Examples showing a large impact of environmental drivers on the speciation process are indeed growing in the literature (Nosil, 2012). However, in more cases than anticipated, the initiation of speciation appears to be linked to non-adaptive intrinsic barriers, even for ecologically divergent species pairs (Presgraves, 2010). Our results show that most genomic regions resisting gene flow between *C. arcania* and *C. gardetta* are not under positive selection, even though signatures of selection, putatively linked to the adaptation of *C. gardetta* to high elevation, can be found in a small proportion of them. The precise contribution of neutral *versus* adaptive processes on the evolution of reproductive isolation is difficult to estimate without knowing the size effect of the genes involved. However, the large proportion of barrier loci with no signal of positive selection suggests that the build-up of reproductive isolation between the ecologically divergent *C. arcania* and *C. gardetta* was largely influenced by intrinsic non-adaptive barriers.

### Divergence with gene flow but mostly isolation

The demographic context of a speciation event, and especially the extent and continuity of gene flow between the diverging populations, can strongly influence the processes initiating or maintaining reproductive isolation. A secondary contact scenario between *C. arcania* and *C. gardetta* appears likely, with putatively strong isolation during the initial stage of the speciation process. The split between the ancestral populations of *C. arcania* and *C. gardetta* would have happened at the beginning of the Pleistocene around 1.8 million years ago, in agreement with previous estimates (Kodandaramaiah & Wahlberg, 2009; Capblancq *et al*., 2015). This geological period is known to be associated with climate cooling and with an increase in speciation events (Levsen *et al*., 2012; April *et al*., 2013; Nevado *et al*., 2018). At that time, a fraction of the species’ ancestral population likely became isolated and gave birth to the current *C. gardetta* lineage. Our results also confirmed the presence of gene flow during the speciation of *C. arcania* and *C. gardetta,* since a secondary contact ∼400,000 generations ago. However, migration between the two species is relatively recent and at low frequency with only a few effective individuals per generation (0.5 – 1.56). While these numbers are comparable with other hybridizing butterfly species with similar divergence time such as *Heliconius melpomene* and *cydno* (Martin *et al*., 2015), they usually correspond to species pairs that have experienced gene flow all along their divergence process, and thus much more cumulative migration throughout their speciation. It is therefore difficult to assess how much migration is associated with effective gene flow between *C. arcania* and *C. gardetta*, especially knowing that mostly F1 hybrids are observed in places where the two species co-occur (Capblancq *et al*., 2019).

The genomics of speciation have been largely studied in pairs of species for which the speciation scenario involves gene flow all along the divergence process (Feder *et al*., 2012). A pure isolation with migration (IM) scenario is nonetheless unlikely for a lot of hybridizing temperate species, which often encompassed period of strict isolation due to large range shifts associated with climate fluctuations (De Jode *et al*., 2023). This *Coenonympha* species pair is one example where migration was interrupted during a large part of the speciation process. In these circumstances, homogenizing gene flow was probably not strong (and/or continuous) enough to lower differentiation at neutral genomic regions in comparison with divergently selected regions, only slightly affecting the genomic landscape of the two species. This pushed us to go beyond the search for contrasts in genetic diversity and differentiation that is widely used in the literature (Turner *et al*., 2005; Nosil *et al*., 2009), to identify the genomic bases of speciation in our species pair.

### Locating the genomic regions contributing to reproductive isolation

The use of process-based procedures was decisive to distinguish the genomic regions involved in reproductive isolation from the regions pervasive to gene flow. The principle behind these methods is relatively simple: scanning the genome of two hybridizing species in search for regions resisting gene flow (Fraïsse *et al*., 2021; Laetsch *et al*., 2023). Recently published procedures use demographically explicit models to test the superiority of a scenario without gene flow over one allowing migration, at the genomic window level. We compared the outcome of two of these methods: DILS, developed by Fraïsse and colleagues (2021) and an adaptation of gIMble, developed by Laetsch and colleagues (2023): the Δ*_B_* method. The two procedures identified multiple regions acting as barriers between *C arcania* and *C. gardetta*, with one large barrier on the Z chromosome and the rest distributed on nine of the 29 autosomes. These were not necessarily located within genomic windows showing extreme patterns of genetic diversity and differentiation. Nonetheless, the results of DILS and Δ*_B_* are not totally concordant, except for the strong signal on the Z chromosome. It could suggest a different sensitivity of the two methods to the overall heterogeneity of gene flow and/or effective population sizes, or variation in their stringency. It is important to note that we analyzed the Z chromosome together with the autosomes here, which is sometimes avoided in the literature (Laetsch *et al*., 2023). While the lower effective population size of the Z chromosome is accounted for in this type of procedure, which re-infer local diversity at the genomic window level, differences in migration rates between the sex chromosomes and the autosomes (e.g., sex-biased migration) could lead to erroneous results. We are not aware of such context in *Coenonympha* butterflies but we need to keep in mind that it could greatly lower the number of barrier loci identified on the Z chromosome (Supplemental Fig. S3).

Exploring the genomic landscape of diversity within and differentiation between species also provided important information. Tajima’s *D*, for example, was low on average but show extreme troughs of negative values at multiple restricted genomic regions. Extremely negative values of Tajima’s *D* imply a recent and strong increase of low frequency alleles, which, when forming a peak in a specific genomic location, is usually linked to a recent selective sweep (Carlson *et al*., 2005). We observed multiple valleys of very low Tajima’s *D* in *C. gardetta* population. They likely originated in response to the selection of alleles allowing the species adaptation to alpine-like environment during the speciation process. Within or close to these Tajima’s *D* troughs we also found signals of recurrent selection, which refers to a scenario where a selective pressure that already acted on the ancestral population still acts on the diverging lineages, reducing polymorphisms within and net divergence between species while increasing differentiation (Han *et al*., 2017; Irwin *et al*., 2018; Shang *et al*., 2021). Therefore, genes important for adaptation, both within and between species, to varying environment are good candidates for recurrent selection. Soft selective sweeps on standing genetic variation (*i.e.*, recurrent selection) were for example found as the main source of positive divergent selection in songbirds (Manthey *et al*., 2021). Some of the regions identified under recurrent selection in *C. arcania* and *C. gardetta* may be under positive selection within and divergent selection between species, which, associated with genetic hitchhiking, can result in genomic blocks of strong differentiation (Charlesworth & Campos, 2014). A more thorough genomic study at the intraspecific level would be necessary to confirm if some of the putative speciation genes identified below could be involved in local adaptation within one or the two species.

### The evolutionary processes underlying the formation of genomic barriers

Our results show that 6.6% of the genome would be acting as barrier to gene flow between *C. arcania* and *C. gardetta.* This is three times more than what was found in a pair of hybridizing *Heliconius* butterflies by Laetsch and colleagues (2023), who show that many loci of weak effect, in addition to 25 strong effect loci, were contributing to reproductive isolation. Another recent study also suggests that reproductive isolation, even in the case of sympatric speciation, could be more polygenic than anticipated (Kautt *et al*., 2020). The barriers identified between the *Coenonympha* species pair host at least 832 protein-coding genes, and many intergenic regions that could also harbor important regulatory regions. While the exact contribution of each genomic region to reproductive isolation (*i.e.*, their effect sizes) is unknown, it still supports a polygenic nature of reproductive isolation in this case as well.

The vast majority of the barriers identified on the autosomes (92.5% of the 291 autosomal barriers) did not show signals of directional positive selection. Resistance to gene flow between *C. arcania* and *C. gardetta* could then be largely driven by non-adaptive processes, suggesting an important contribution of endogenous barriers to the evolution of reproductive isolation in this species pair. Given that the first 1.5 Myrs of the speciation process was likely realized in strong isolation between the diverging lineages, we can hypothesize that many genes and/or regulatory regions accumulated independent mutations in *C. arcania* and *C. gardetta* genomes. This is supported by the very high genetic differentiation observed on average between the two species. The recombination of co-evolved alleles in hybrids is very often associated with negative interactions, leading to reduced hybrid fitness and increased isolation between the diverging lineages (Coyne, 1992). These endogenous barriers could have arisen as a major driver of reproductive isolation between *C. arcania* and *C. gardetta* during the initial stage of their divergence, while they were not exchanging genetic material. However, adaptive signals were still found in a small proportion of the autosomal barriers, suggesting some influence of exogeneous barriers in the evolution of reproductive isolation. Because selection was supposedly already acting on these genomic regions before the divergence (*i.e.*, recurrent selection), we can hypothesize that local adaptations in the ancestral population are now differentiating the two species and participate to their reproductive isolation. That would fit well with the strong ecological divergence observed between *C. arcania* and *C. gardetta*, and the identification of many genes involved in adaptation to hypoxia in the barriers showing signals of positive selection. Selection against hybrids would then be partially linked to environmental filtering of unfit phenotypes, reinforcing post-zygotic isolation between the two species. Still on the autosomes, many windows subject to positive selection were found in genomic regions permeable to gene flow (56 windows outside barriers and 22 inside), confirming the potential of our framework to differentiate positive selection contributing or not to reproductive isolation (Laetsch *et al*., 2023).

The case of the Z chromosome is striking but also delicate to interpret. We identified the Z chromosome as homogeneously resistant to gene flow, supporting that the Z chromosome is a large endogenous and/or exogeneous barrier to reproduction between *C. arcania* and *C. gardetta*. Unlike on the autosomes, the patterns of genetic diversity and differentiation were not contrasted enough on the Z chromosome to easily differentiate the influence of adaptive and non-adaptive processes on shaping the genomic landscape. The proportion of genes with positive DoS (40%) is relatively important on the Z chromosome but most genes still return negative DoS values, indicating that both positive and purifying selection are at play. We also did not find any molecular pathway particularly enriched on the Z chromosome or known important adaptive genes. An entire chromosome the size of the Z chromosome (25.8 Mbp) responding homogeneously to positive selection is also rather unlikely. In that regard, the accumulation of intrinsic non-adaptive genetic incompatibilities could appear as the most likely contributor of reproductive isolation on the Z chromosome as well. Overall, an important role of intrinsic genetic incompatibilities in *C. arcania* and *C. gardetta* reproductive isolation agrees with the results of experimental interspecific crosses that found that F1 hybrids were mostly sterile or very unfit (de Lesse, 1960).

### A large Z-effect probably at play in Coenonympha

Sex chromosomes are known to play an exacerbated role in the divergence of many sexual organisms (Presgraves, 2018) and Lepidoptera in particular (Sperling, 1994). A “large X(or Z)-effect” correspond to an exacerbated divergence of the X (or Z) chromosome produced by selection against unfit or sterile hybrids (Coyne, 1992). It assumes the accumulation of incompatible alleles and the involvement of the sex chromosomes in post-zygotic barriers to gene flow between diverging populations or species (Presgraves, 2002). Our results support that a large Z-effect is at play in the divergence process of *C. arcania* and *C. gardetta*. First, we found the almost entire Z chromosome acting as a barrier between *C. arcania* and *C. gardetta*, confirming its preponderant role in the evolution of reproductive isolation. Second, we found extreme differences in genomic diversity and differentiation between the autosomes and the Z chromosome, which can indicate an exacerbated influence of the sex chromosomes (Presgraves, 2018; Battey, 2020). The ratio *F*_ST(Z)_ / *F*_ST(A)_ = 1.92 is much higher in the studied *Coenonympha* species than for most butterfly species pairs (1.44) (Presgraves, 2018). On the contrary, the π_(Z)_/π_(A)_ is extremely low for both *C. arcania* (0.51) and *C. gardetta* (0.44). The effective population size of the Z chromosome is ¾ of the one of an autosome, leading to a theoretical diversity ratio of π_(Z)_/π_(A)_ ∼ 0.75 (Charlesworth, 2001), much higher than the one we observed. Unbalanced fertility between sex and/or age classes can lower down this ratio but theory predicts that values below 0.64 would most likely imply the involvement of selection (Charlesworth, 2001). Recent bottlenecks can also influence the π_(Z)_/π_(A)_ ratio (Pool & Nielsen, 2007) but we inferred very large effective population sizes for both species, making a recent and strong bottleneck unlikely as the driver of reduced diversity on the Z chromosome.

### Adaptive genes between C. arcania and C. gardetta

In the barriers under positive selection, we identified multiple genes potentially associated with response to hypoxia and other stresses. Those would be consistent with the hypothesis of adaptation to high altitude in *C. gardetta,* which involves adapting to lower oxygen pressures and more frequent stresses. For example, the serine/threonine protein kinase, coded by a gene on chromosome 21, is suggested to be linked to activation of mitochondrial respiration in response to a decrease in ATP levels, which happens after a stress and particularly during hypoxia (Murray, 2009). We also found one gene coding for an apoptosis-inducing factor on chromosome 9. Apoptosis-inducing factors are known to be involved in adaptation to high elevation in humans (Sharma *et al*., 2022) and mammals (Wu *et al*., 2020), due to their role in regulating apoptosis of the cells during hypoxia. In the same lines, two phosphatidylserine synthases, which act as recognition receptor in apoptotic cells (Naeini *et al*., 2020) were annotated on chromosome 3, in one of the most interesting genomic barriers. Furthermore, one gene producing a fatty acid synthetase was found on chromosome 15, and a reduced oxidation of fatty acids is suggested to be beneficial at high elevation when hypoxia makes oxidation of carbohydrates more favorable (Ge *et al*., 2012). Those genes are located on various autosomes and the Z chromosome, suggesting that such adaptation to elevation in *C. gardetta* would be polygenic. The barrier loci also host genes involved in cell functioning, organism development and DNA or RNA repair and expression. Although there is no certainty about the implication of these genes in the speciation process, they might be involved in genomic incompatibilities at the developmental stage or hybrid infertility (Palopoli & Wu, 1994), which would explain the lack of backcrossed individuals found in the field even though F1 hybrids are present. The usual suspects such as *cortex*, *optix* or *WntA*, which are genes classically involved in butterfly speciation (Van Belleghem *et al*., 2021), were found nowhere near our identified genomic barriers to gene flow. That could suggest a weak role of wing coloration, patterning and shape in the isolation of this species pair, contrary to many other examples of tropical and temperate butterflies (Jiggins, 2006; Mavárez *et al*., 2006; Salazar *et al*., 2010). Important regulatory regions or structural variants could be involved as well in the reproductive isolation between *C. arcania* and *C. gardetta*. Improving our understanding of the *Coenonympha* genome and its functional annotation in the coming years will be an important step to look beyond the genes and integrate more complex genetic features likely involved in the speciation process.

## MATERIALS & METHODS

### Genomic data acquisition

Nineteen individuals of *Coenonympha arcania* and 17 individuals of *C. gardetta* were sampled across the two species ranges in Europe, with a particular focus on the Alps and their surroundings (Supplemental Table S2). The body of each individual was stored in 90% alcohol and kept at −20 °C until processing. Whole genomic DNA was extracted in the lab from the thorax of each sample using the Qiagen DNeasy blood and tissue kit. Extracted DNA was sent to the Genotoul platform (https://www.genotoul.fr) for whole genome sequencing library preparation using the TruSeq Nano DNA Illumina’s Library Preparation kit. The 37 libraries were then sequenced on a lane of Illumina NovaSeq 6000 S4 to generate paired-end 150-bp reads. Read sequences were demultiplexed, clipped of Illumina adapter sequences, and trimmed of low quality flanking sequence (Q<20) using a sliding window of 6 bp using Trimmomatic (Bolger *et al*., 2014). Mapping of the sequence reads was performed with the program BWA (Li & Durbin, 2009), using the BWA-MEM algorithm. As a reference, we used the new BST1 assembly of *C. arcania* genome described in (Legeai *et al*., 2024). The SAM files resulting from the aligned reads were converted to BAM files using SAMtools (Li *et al*., 2009). PCR duplicates were removed using the *“markdup”* function in sambamba-0.6.8 (Tarasov *et al*., 2017). The filtered BAM files were sorted and indexed using SAMtools.

We used ANGSD (Analysis of Next Generation Sequencing Data) (Korneliussen *et al*., 2014), to produce genotype likelihoods for each individual and covered genomic site. ANGSD was first ran independently for each species using the SAMtools genotype likelihood model, only using reads having unique best hits, setting a minimum MapQ score to keep a read to 20, a min nucleotide Q score to consider a site to 20, a minimum number of 5 individuals with coverage to consider a site, a minimum of 3 and maximum of 60 reads to estimate genotype likelihood for one individual, keeping only biallelic sites, performing the base alignment quality (BAQ: Phred-scaled probability of a read base being misaligned)(Li, 2011) as in SAMtools. We then identified the genomic sites (monomorphic and polymorphic) that were covered for both species and re-ran ANGSD with all samples using the “-site” option to produce a global dataset. For downstream analyses, we either directly used the genotype likelihoods or “hard called” genotypes using a posterior probability cutoff of 80% (more details in Supplemental Code S1).

### Inference of historic population dynamics and species divergence

A Site Frequency Spectrum (SFS) was inferred and optimized for each species from the genotype likelihoods using the realSFS sub-program of ANGSD. A total sequence length of 203 Mb was used for creating the SFS, including monomorphic and polymorphic sites, all covered in both species. We used Fastsimcoal2 (Excoffier *et al*., 2013) to compare the likelihood of different speciation scenarios, including strict isolation (SI), isolation with migration (IM), secondary contact (SC) and ancient migration (AM). To avoid making the models overly complex, we modeled gene flow as homogeneous along the genomes and both gene flow and *N_e_* as homogeneous through time for a specific period. These models were compared running 30 independent maximizations of the likelihood based on the observed joint SFS derived from the two species SFS, and retaining the run/model with the lowest AIC. Once the most likely scenario was identified, we estimated the timing of divergence (*T*_split_), the timing of gene flow (*T*_gene-flow_), the current populations sizes (*N*_arcania_, *N*_gardetta_), and the migration rate between species (*m*_arcania-gardetta_, *m*_gardetta-arcania_). These parameters were estimated running 30 independent maximizations of the likelihood based on the observed joint SFS derived from the two species SFS, and retaining the estimate with the highest likelihood. We then performed 100 parametric bootstraps to obtain the 95% confidence interval for each parameter estimate.

In parallel, we used the procedure implemented in DILS (Fraïsse *et al*., 2021) to compare the same divergence models with an ABC framework. DILS was run on 9,840 sets of sequences representing the genotype data for the 36 samples on 20 kbp long genomic windows along the entire reference genome. For a random selection of 1000 sequences, DILS first conducts a genome-wide analysis using an ABC procedure based on random forest and a large set of summary statistics. This first step of the procedure identifies the genome-wide model, and associated parameters, that reproduce best the observed dataset. We checked the validity of the procedure using goodness-of-fit tests (Supplemental Table S3 and Fig. S4). Genetic data were simulated under the best supported model, using the estimated parameters. These simulations then empirically produce the statistical distributions summarized under the inferred model. We then examine whether the value observed from the *Coenonympha* dataset is correctly captured by the inferred model for each of the summary statistics.

### Locating barrier loci in the genomes

To look for barrier loci between *C. arcania* and *C. gardetta*, we tested a local reduction of gene flow along the genome using an adaptation of the model-based approach developed in gIMble (Laetsch *et al*., 2023) and the procedure implemented in DILS (Fraïsse *et al*., 2021). We had to modify the available version of gIMble because it assumes an isolation with migration scenario between the diverging populations when the most likely scenario of speciation for *C. arcania* and *C. gardetta* was a secondary contact after a period of strict isolation (see Results). Otherwise, we followed a similar procedure, which tests if the migration rate estimated for a specific genomic region (*m_e,i_*) is reduced compared to the migration rate inferred for the entire genome (m□*_e_*), using coalescent modeling. In our case, the likelihood of a secondary contact (SC) model was compared to a model of strict isolation (SI) for each genomic region of interest. In the first model of divergence, *m_e,i_* is fixed to m□*_e_*, the divergence time (*T_split_*) and the timing of secondary contact (*T_sc_*) are also fixed to their genome-wide estimates and only the effective population sizes *N_arcania_* and *N_gardetta_* are inferred to account for heterogeneity in genetic diversity along the genome. In the second model *m*_e_ is fixed to 0 (SI) while *N_arcania_* and *N_gardetta_* are inferred, and divergence time (*T_split_*) is fixed to its genome-wide estimates. Finally, the difference in likelihood (Δ*_B_*) between the two models is estimated to determine if having *m_e,i_* = 0 improves the fit of the model to the data. The genomic regions for which Δ*_B_* > 0 show a reduced local migration rate compared to the genome-wide estimate and can be considered as putatively involved in reproductive isolation. To apply that procedure on our data, we first used realSFS to estimate and optimize a local 2D-SFS between *C. arcania* and *C. gardetta* for each 50kb genomic window described above. We then used Fastsimcoal2 (Excoffier *et al*., 2013) to estimate, for each window, the likelihood and parameters of the two models described above. We estimated Δ*_B_* for each genomic window and qualified a region as barrier to gene flow when Δ*_B_* > 0. To account for a possible difference in migration rates between sex chromosomes and autosomes we re-ran the procedure independently for the Z chromosome, using an overall migration rate estimated with this chromosome only. However, if the Z chromosome hosted many barriers, this Z focused procedure would lead to underestimating the real migration rate and therefore the number of barriers on this chromosome. We acknowledge that this is rather circular and unfortunately impossible to resolve precisely without knowing the nature and frequency of migration events between species. Finally, uncertainty in the procedure was quantified by estimating a false positive rate (FPR) for each window. We simulated 100 replicates under the locally best fitting set of N estimates but fixing *m_e_* to m□*_e_* and *T_split_* to the time estimate found with the global model. We then used the same procedure as above to measure the fraction of the simulation replicates for which ΔB > 0.

We compared the Δ*_B_* results to the outputs of DILS, which, in addition to inferring demographic parameters, look for barrier loci in the genomes (Fraïsse *et al*., 2021). DILS was executed with the same input file that was described in the “*demographic inferences*” section. The random forest-based ABC procedure was conducted on each genomic window independently to decide between a scenario of divergence with secondary contact (SC) and a scenario of strict isolation (SI) at the 50kb window level, while accounting for potential variation in effective population size across genomic regions. The mode “*bimodal”* was used (two categories of locus in the genome: proportion *P* linked to barriers with migration rate of zero, 1-*P* not linked to barriers with non-null migration) and we used a mutation rate of 2.9×10^−9^ for the simulations. Windows returning a SI posterior probability > 0.7 were considered as barrier loci.

### Species genomic diversity along the genome

To characterize patterns of genetic diversity along BST1 reference genome for the two species, we first estimated two population-specific parameters with Tajima’s *D* (Tajima, 1989) and pairwise nucleotide diversity (π). We measured these parameters for each species independently using the same filtering options, as described above. Pairwise diversity (π) and Tajima’s *D* were calculated on abutting genomic windows of 50kb using the realSFS subprogram of ANGSD and its function “*thetaStat*” (Korneliussen *et al*., 2013) from the species-specific genotype likelihood files. To look at the heterogeneity of genetic divergence along the genome of the two species, we estimated *d*_xy_, the absolute genetic divergence, *d*_a_, the net genetic divergence, and *F*_ST_, as a proxy of genetic differentiation, using the same 50kb abutting genomic windows as for the species-specific parameters described above. *d*_xy_ was estimated following Toews et al. (2016). For each polymorphic site we measured the probability of sampling different alleles between the two species using the species-specific ANGSD allele frequency files (.mafs files) and the formula: *d_xv_* = *f*_1_ * (1-*f*_2_) + *f*_2_ * (1-*f*_1_), where *f*_1_ is the minor allele frequency in species 1 and *f*_2_ is the minor allele frequency in species 2. The site-specific estimates obtained were then averaged within each genomic window, including monomorphic sites as 0s. Net divergence *d*_a_ was estimated for each genomic window as follow: 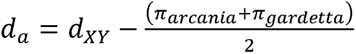. Mean pairwise *F*_ST_ values were also estimated for each window from the optimized SFS produced for each species as described above and using the function “*fst*” of the realSFS program. Pairwise LD was estimated using the program ngsLD (Fox *et al*., 2019), restricting comparisons to pairs of polymorphic sites within each window of 50 kb used above for the estimation of the genetic diversity parameters. LD decay was estimated for each genomic window by weighting the *r*^2^ of the correlation between loci by their physical distance (bp) along the genome. The averaged value was recorded for each window and used as a proxy of linkage strength along the genome.

### Direction of Selection (DoS)

We used SNPeff (Cingolani *et al*., 2012) and the positions along the *BST1* reference genome to annotate the variants to functional classes: upstream and downstream of genes, introns, synonymous, nonsynonymous, intronic, or intergenic sites. We then used this information and the frequency of the different alleles within *C. arcania* and *C. gardetta* populations to estimate, for all protein-coding genes, the direction of selection (DoS) statistics proposed in (Stoletzki & Eyre-Walker, 2011). This metric results from the adaptation of the McDonald-Kreitmand (MK) test for selection (McDonald & Kreitman, 1991), where the number of synonymous and nonsynonymous substitution (*D*_n_ and *D*_s_) are compared to the number of synonymous and nonsynonymous polymorphisms (*P*_n_ and *P*_s_). The direction of selection is estimated as follows: DoS = *D*_n_ / (*D*_n_ + *D*_s_) - *P*_n_ / (*P*_n_ + *P*_s_), with DoS > 0 suggesting positive selection and DoS < 0 suggesting purifying selection on deleterious mutations (Stoletzki & Eyre-Walker, 2011). In our case *D*_n_ and *D*_s_ were mutations with divergently fixed alleles in *C. arcania* and *C. gardetta* populations when *P*_n_ and *P*_s_ represented mutations that were polymorphic in one or the two species.

### Signals of selection along the genomes

To locate genomic windows under selection we followed the “correlation of genomic landscapes” approach proposed in Shang et al. (2021) and adapted form Han et al. (2017) and Irwin et al. (2018), which differentiates four evolutionary scenarios based on extremeness of genomic parameters. Each scenario corresponds to specific divergence and selection history with each history being characterized by different patterns of genetic differentiation (*F*_ST_), net divergence (*d*_XY_) and nucleotide diversity (π). Divergence with gene flow, when selection acts only at specific genes involved in reproductive isolation while gene flow is pervasive elsewhere in the genome, results in high *F*_ST_, high *d*_XY_ and low π. Allopatric selection, when selection acts independently within the two isolated species, results in high *F*_ST_, average *d*_XY_ and low π. Recurrent selection, which relates to similar selective pressure in the two diverging lineages and their ancestral population, results in high *F*_ST_, low *d*_XY_ and low π. Balancing selection, where ancestral polymorphism is maintained by selection, results in low *F*_ST_, high *d*_XY_ and high π. We considered the parameters as being “low” or “high” when they were in bottom or top 5% of the distribution, respectively, and “average” when they fell between the 30^st^ and 70^st^ percentiles. This characterization was conducted independently on the Z chromosome and the autosomes.

### Exploration of the candidate genes

Four types of genomic windows were characterized based on their association with a barrier locus (“linked to barrier” or “unlinked to barrier”) and if they showed signals of positive selection (divergent, recurrent or allopatric) with the “correlation of genomic landscapes” approach described above (“positive selection” or “no positive selection”). To explore further the potential pathways involved in reproductive isolation between *C. arcania* and *C. gardetta* we then looked at the genes located in these four types of genomic regions. To assess if these lists of annotated genes were overrepresented for different molecular functions or biological processes, we conducted for each group of windows a gene ontology (GO) term enrichment analysis using the *topGO* R-package (Alexa & Rahnenfuhrer, 2023) and tested the significance of the enrichment using a Kolmogorov Smirnov test. Finally, we exhaustively looked at the genes located in windows linked to barrier and showing signals of positive selection.

## DATA ACCESS

Sequence data generated in this study have been submitted to the NCBI BioProject database (https://www.ncbi.nlm.nih.gov/bioproject/1053281), under accession number PRJNA1053281.

Custom scripts to reproduce the analyses conducted in the study can be found in the Supplemental Code S1 and S2, as well as on Github: https://github.com/Capblancq/Speciation-Coenonympha-butterflies/tree/master/Speciation-genomics-arcania-gardetta

## ACKNOWLEDGEMENTS

This project was supported by the French National Research Agency (ANR-20-CE02-0017). We appreciate the assistance of Jesus Mavarez and Paul Doniol-Valcroze for their contribution to the collecting of material used for the genomic study and of Christelle Fraïsse for interesting discussions on our results.

## AUTHOR CONTRIBUTIONS

Laurence Després, Mathieu Joron and Thibaut Capblancq conceived the study and collected the samples. TC carried out the laboratory work, analyzed genomic data with the help of Fabrice Legeai and conducted the statistical analyzes with help from Camille Roux and Frédéric Boyer. TC wrote a first draft of the manuscript and all authors provided critical feedback.

## REFERENCES

Alexa A, Rahnenfuhrer J. 2023. topGO: Enrichment Analysis for Gene Ontology.

April J, Hanner RH, Dion-Côté A-M, Bernatchez L. 2013. Glacial cycles as an allopatric speciation pump in north-eastern American freshwater fishes. Molecular Ecology 22: 409–422.

Barton NH, Hewitt GM. 1985. Analysis of hybrid zones. Annual Review of Ecology and Systematics 16: 113–148.

Bateson W. 1909. Heredity and variation in modern lights. Darwin and modern science.

Battey CJ. 2020. Evidence of linked selection on the Z chromosome of hybridizing hummingbirds*. Evolution 74: 725–739.

Bolger AM, Lohse M, Usadel B. 2014. Trimmomatic: A flexible trimmer for Illumina sequence data. Bioinformatics 30: 2114–2120.

Capblancq T. 2016. La spéciation hybride□: réflexions générales et exploration d ‘ un cas d’ étude chez des papillons alpins du genre Coenonympha.

Capblancq T, Després L, Rioux D, Mavárez J. 2015. Hybridization promotes speciation in Coenonympha butterflies. Molecular Ecology 24: 6209–6222.

Capblancq T, Mavárez J, Rioux D, Després L. 2019. Speciation with gene flow: Evidence from a complex of alpine butterflies (Coenonympha, Satyridae). Ecology and Evolution 9: 6444–6457.

Carlson CS, Thomas DJ, Eberle MA, Swanson JE, Livingston RJ, Rieder MJ, Nickerson DA. 2005. Genomic regions exhibiting positive selection identified from dense genotype data. Genome Research 15: 1553–1565.

Charlesworth B. 2001. The effect of life-history and mode of inheritance on neutral genetic variability. Genetical Research 77: 153–166.

Charlesworth B. 2012. The Effects of Deleterious Mutations on Evolution at Linked Sites. Genetics 190: 5–22.

Charlesworth B, Campos JL. 2014. The Relations Between Recombination Rate and Patterns of Molecular Variation and Evolution in *Drosophila*. Annual Review of Genetics 48: 383–403.

Charlesworth B, Nordborg M, Charlesworth D. 1997. The effects of local selection, balanced polymorphism and background selection on equilibrium patterns of genetic diversity in subdivided populations. Genetical Research 70: 155–174.

Cingolani P, Platts A, Wang LL, Coon M, Nguyen T, Wang L, Land SJ, Lu X, Ruden DM. 2012. A program for annotating and predicting the effects of single nucleotide polymorphisms, SnpEff. Fly 6: 80–92.

Coyne JA. 1992. Genetics and speciation. Nature 355: 511–515.

Coyne JA, Orr HA. 2004. Speciation. Sinauer associates Sunderland, MA.

Cruickshank TE, Hahn MW. 2014. Reanalysis suggests that genomic islands of speciation are due to reduced diversity, not reduced gene flow. Molecular Ecology 23: 3133–3157.

De Jode A, Le Moan A, Johannesson K, Faria R, Stankowski S, Westram AM, Butlin RK, Rafajlović M, Fraïsse C. 2023. Ten years of demographic modelling of divergence and speciation in the sea. Evolutionary Applications 16: 542–559.

de Lesse. 1960. Zoologie et Biologie Animale. Paris.

Dobzhansky T. 1936. Studies on Hybrid Sterility. II. Localization of Sterility Factors in Drosophila Pseudoobscura Hybrids. Genetics 21: 113–135.

Endler JA. 1977. Geographic variation, speciation, and clines. Princeton University Press.

Excoffier L, Dupanloup I, Huerta-Sánchez E, Sousa VC, Foll M. 2013. Robust Demographic Inference from Genomic and SNP Data. PLoS Genetics 9.

Feder JL, Egan SP, Nosil P. 2012. The genomics of speciation-with-gene-flow. Trends in Genetics 28: 342–350.

Fox EA, Wright AE, Fumagalli M, Vieira FG. 2019. ngsLD: evaluating linkage disequilibrium using genotype likelihoods. Bioinformatics: 1–2.

Fraïsse C, Popovic I, Mazoyer C, Spataro B, Delmotte S, Romiguier J, Loire É, Simon A, Galtier N, Duret L, et al. 2021. DILS: Demographic inferences with linked selection by using ABC. Molecular Ecology Resources 21: 2629–2644.

Ge R-L, Simonson TS, Cooksey RC, Tanna U, Qin G, Huff CD, Witherspoon DJ, Xing J, Zhengzhong B, Prchal JT, et al. 2012. Metabolic insight into mechanisms of high-altitude adaptation in Tibetans. Molecular Genetics and Metabolism 106: 244–247.

Han F, Lamichhaney S, Grant BR, Grant PR, Andersson L, Webster MT. 2017. Gene flow, ancient polymorphism, and ecological adaptation shape the genomic landscape of divergence among Darwin’s finches. Genome Research 27: 1004–1015.

Irwin DE, Milá B, Toews DPL, Brelsford A, Kenyon HL, Porter AN, Grossen C, Delmore KE, Alcaide M, Irwin JH. 2018. A comparison of genomic islands of differentiation across three young avian species pairs. Molecular Ecology 27: 4839–4855.

Jiggins CD. 2006. Speciation: Reinforced butterfly speciation. Heredity 96: 107–108.

Kautt AF, Kratochwil CF, Nater A, Machado-Schiaffino G, Olave M, Henning F, Torres-Dowdall J, Härer A, Hulsey CD, Franchini P, et al. 2020. Contrasting signatures of genomic divergence during sympatric speciation. Nature 588: 106–111.

Kodandaramaiah U, Wahlberg N. 2009. Phylogeny and biogeography of Coenonympha butterflies (Nymphalidae: Satyrinae)–patterns of colonization in the Holarctic. Systematic Entomology 34: 315–323.

Korneliussen TS, Albrechtsen A, Nielsen R. 2014. ANGSD: Analysis of Next Generation Sequencing Data. BMC Bioinformatics 15: 356–368.

Korneliussen TS, Moltke I, Albrechtsen A, Nielsen R. 2013. Calculation of Tajima’s D and other neutrality test statistics from low depth next-generation sequencing data. BMC Bioinformatics 14.

Laetsch DR, Bisschop G, Martin SH, Aeschbacher S, Setter D, Lohse K. 2023. Demographically explicit scans for barriers to gene flow using gIMble (N Bierne, Ed.). PLOS Genetics 19: e1010999.

Legeai F, Romain S, Capblancq T, Doniol-Valcroze P, Joron M, Lemaitre C, Després L. 2024. Chromosome-Level Assembly and Annotation of the Pearly Heath *Coenonympha arcania* Butterfly Genome (C Wheat, Ed.). Genome Biology and Evolution 16: evae055.

Levsen ND, Tiffin P, Olson MS. 2012. Pleistocene Speciation in the Genus Populus (Salicaceae). Systematic Biology 61: 401.

Li H. 2011. Improving SNP discovery by base alignment quality. Bioinformatics 27: 1157–1158.

Li H, Durbin R. 2009. Fast and accurate long-read alignment with Burrows-Wheeler transform. Bioinformatics 25: 1754–1760.

Li H, Handsaker B, Wysoker A, Fennell T, Ruan J, Homer N, Marth G, Abecasis G, Durbin R. 2009. The Sequence Alignment/Map format and SAMtools. Bioinformatics 25: 2078–2079.

Manthey JD, Klicka J, Spellman GM. 2021. The Genomic Signature of Allopatric Speciation in a Songbird Is Shaped by Genome Architecture (Aves: *Certhia americana*) (K Lohmueller, Ed.). Genome Biology and Evolution 13: evab120.

Martin SH, Dasmahapatra KK, Nadeau NJ, Salazar C, Walters JR, Simpson F, Blaxter M, Manica A, Mallet J, Jiggins CD. 2013. Genome-wide evidence for speciation with gene flow in *Heliconius* butterflies. Genome Research 23: 1817–1828.

Martin SH, Eriksson A, Kozak KM, Manica A, Jiggins CD. 2015. Speciation in Heliconius Butterflies: Minimal Contact Followed by Millions of Generations of Hybridisation. Evolutionary Biology.

Mavárez J, Salazar CA, Bermingham E, Salcedo C, Jiggins CD, Linares M. 2006. Speciation by hybridization in Heliconius butterflies. Nature 441: 868–871.

McDonald J H, Kreitman M. 1991. Adaptive protein evolution at the Adh locusin Drosophila. Nature 351: 652–654.

Moreira LR, Klicka J, Smith BT. 2023. Demography and linked selection interact to shape the genomic landscape of codistributed woodpeckers during the Ice Age. Molecular Ecology 32: 1739–1759.

Muller HJ. 1942. Isolating mechanisms, evolution, and temperature. In: Biol. Symp. 71.

Murray AJ. 2009. Metabolic adaptation of skeletal muscle to high altitude hypoxia: how new technologies could resolve the controversies. Genome Medicine 1: 117.

Naeini MB, Bianconi V, Pirro M, Sahebkar A. 2020. The role of phosphatidylserine recognition receptors in multiple biological functions. Cellular & Molecular Biology Letters 25: 23.

Nevado B, Contreras-Ortiz N, Hughes C, Filatov DA. 2018. Pleistocene glacial cycles drive isolation, gene flow and speciation in the high-elevation Andes. New Phytologist 219: 779–793.

Noor MAF, Bennett SM. 2009. Islands of speciation or mirages in the desert? Examining the role of restricted recombination in maintaining species. Heredity 103: 439–444.

Nosil P. 2012. Ecological Speciation. Oxford Univiversity Press.

Nosil P, Funk DJ, Ortiz-Barrientos D. 2009. Divergent selection and heterogeneous genomic divergence. Molecular Ecology 18: 375–402.

Palopoli MF, Wu C-I. 1994. Genetics of Hybrid Male Sterility Between Drosophila Sibling Species: A Complex Web of Epistasis Is Revealed in Interspecific Studies. Genetics: 329–341.

Pool JE, Nielsen R. 2007. POPULATION SIZE CHANGES RESHAPE GENOMIC PATTERNS OF DIVERSITY. Evolution 61: 3001–3006.

Presgraves DC. 2002. PATTERNS OF POSTZYGOTIC ISOLATION IN LEPIDOPTERA. Evolution 56: 1168–1183.

Presgraves DC. 2010. The molecular evolutionary basis of species formation. Nature Reviews Genetics 11: 175–180.

Presgraves DC. 2018. Evaluating genomic signatures of “the large X-effect” during complex speciation. Molecular Ecology 27: 3822–3830.

Ravinet M, Faria R, Butlin RK, Galindo J, Bierne N, Rafajlović M, Noor MAF, Mehlig B, Westram AM. 2017. Interpreting the genomic landscape of speciation: a road map for finding barriers to gene flow. Journal of Evolutionary Biology 30: 1450–1477.

Roux C, Fraïsse C, Romiguier J, Anciaux Y, Galtier N, Bierne N. 2016. Shedding Light on the Grey Zone of Speciation along a Continuum of Genomic Divergence. PLoS Biology 14: 1–23.

Salazar C, Baxter SW, Pardo-Diaz C, Wu G, Surridge A, Linares M, Bermingham E, Jiggins CD. 2010. Genetic Evidence for Hybrid Trait Speciation in Heliconius Butterflies. PLoS Genetics 6: e1000930.

Schluter D. 2009. Evidence for Ecological Speciation and Its Alternative. Science 323: 737–741.

Seehausen O, Butlin RK, Keller I, Wagner CE, Boughman JW, Hohenlohe PA, Peichel CL, Saetre G-P, Bank C, Brännström Å, et al. 2014. Genomics and the origin of species. Nature Reviews Genetics 15: 176–192.

Shang H, Rendón-Anaya M, Paun O, Field DL, Hess J, Vogl C, Liu J, Ingvarsson PK, Lexer C, Leroy T. 2021. *Drivers of genomic landscapes of differentiation across* Populus *divergence gradient*. Evolutionary Biology.

Sharma V, Varshney R, Sethy NK. 2022. Human adaptation to high altitude: a review of convergence between genomic and proteomic signatures. Human Genomics 16: 21.

Simon A, Bierne N, Welch JJ. 2018. Coadapted genomes and selection on hybrids: Fisher’s geometric model explains a variety of empirical patterns. Evolution Letters 2: 472–498.

Smadja CM, Butlin RK. 2011. A framework for comparing processes of speciation in the presence of gene flow: FRAMEWORK FOR COMPARING PROCESSES OF SPECIATION. Molecular Ecology 20: 5123–5140.

Sperling FAH. 1994. SEX-LINKED GENES AND SPECIES DIFFERENCES IN LEPIDOPTERA. The Canadian Entomologist 126: 807–818.

Stankowski S, Ravinet M. 2021. Defining the speciation continuum. Evolution 75: 1256–1273.

Stoletzki N, Eyre-Walker A. 2011. Estimation of the Neutrality Index. Molecular Biology and Evolution 28: 63–70.

Tajima F. 1989. Statistical Method for Testing the Neutral Mutation Hypothesis by DNA Polymorphism. Genetics Society of America: 585–595.

Tarasov A, Vilella AJ, Cuppen E, Nijman IJ, Prins P. 2017. Genome analysis Sambamba□: fast processing of NGS alignment formats. Bioinformatics 31: 2032–2034.

Toews DPL, Taylor SA, Vallender R, Brelsford A, Butcher BG, Messer PW, Lovette IJ. 2016. Plumage Genes and Little Else Distinguish the Genomes of Hybridizing Warblers. Current Biology 26: 2313–2318.

Turner TL, Hahn MW, Nuzhdin SV. 2005. Genomic Islands of Speciation in Anopheles gambiae (N Barton, Ed.). PLoS Biology 3: e285.

Van Belleghem SM, Cole JM, Montejo□Kovacevich G, Bacquet CN, McMillan WO, Papa R, Counterman BA. 2021. Selection and isolation define a heterogeneous divergence landscape between hybridizing *Heliconius* butterflies. Evolution 75: 2251–2268.

Wu D-D, Yang C-P, Wang M-S, Dong K-Z, Yan D-W, Hao Z-Q, Fan S-Q, Chu S-Z, Shen Q-S, Jiang L-P, et al. 2020. Convergent genomic signatures of high-altitude adaptation among domestic mammals. National Science Review 7: 952–963.

