## Supplemental Materials for "Untangling the contribution of adaptive *versus* non-adaptive processes in the evolution of reproductive isolation between *Coenonympha* butterflies"

Supplemental Figure S1:

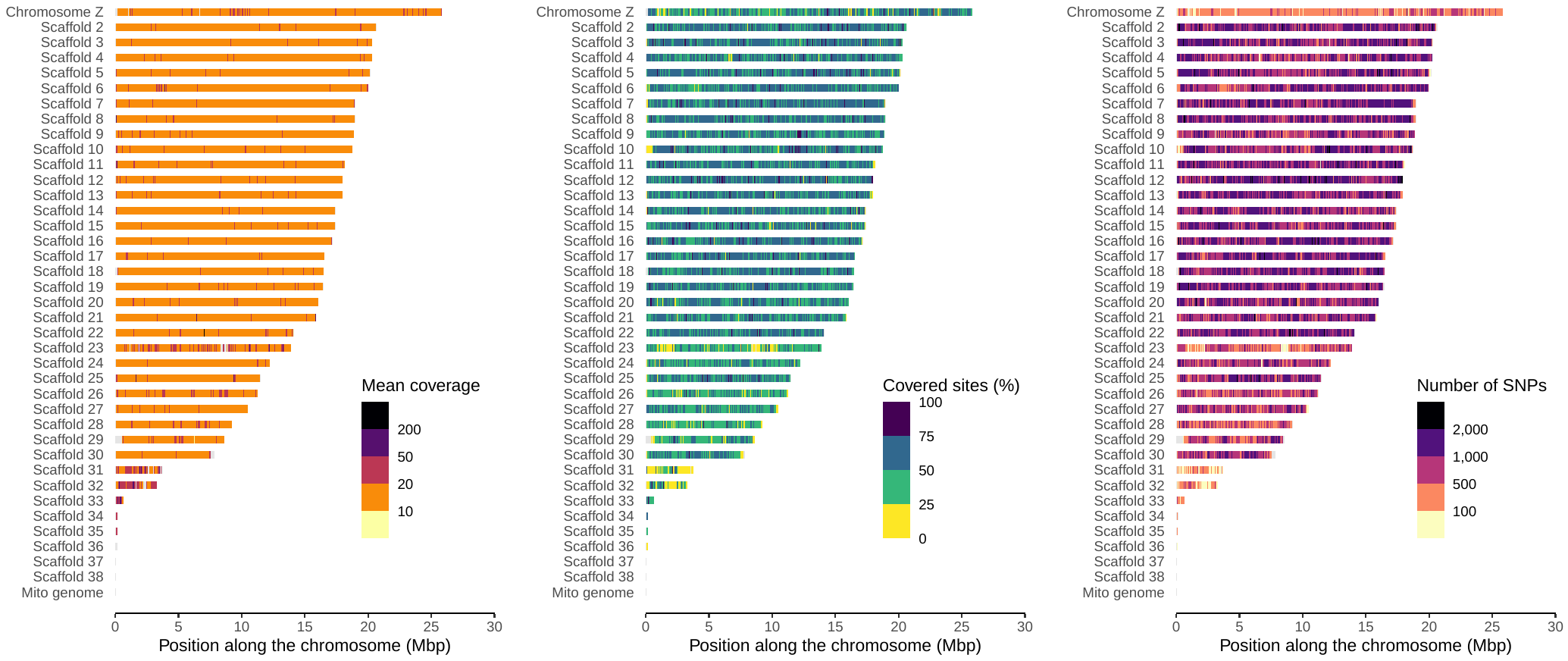

**Fig. S1:** Distribution of sequence coverage and number of polymorphic sites obtained from whole genome resequencing of 36 *Coenonympha* specimens mapped onto the *Coenonympha arcania* BST1 reference genome. The three parameters shown (mean depth of coverage with a mapping quality (MapQ) superior to 20, mean number of sites covered more than 3 times per sample, and mean number of polymorphic sites (SNPs)) were estimated for each of 5,334 abutting windows of 100kb. Grey sections correspond to windows that have not passed the filtering steps and were not used for downstream analyses.

Supplemental Table S1:

To test the potential influence of the Z chromosome on our results, we carried out the same demographic inferences using a 2D-SFS estimated from the autosomes only or including both the autosomes and the Z chromosome or only including the Z chromosome:

| **Fastsimcoal2** | | Including Z chromosome | Excluding Z chromosome | Only Z chromosome |
| --- | --- | --- | --- | --- |
| Best model | | Secondary Contact (SC) | Secondary Contact (SC) | Secondary Contact (SC) |
| Divergence time | |  |  |  |
|  | Tsplit | 1,793,459 | 1,776,965 | *Fixed to whole genome value* |
|  | Tsc | 360,240 | 310,970 | *Fixed to whole genome value* |
| Effective sizes | |  |  |  |
|  | Narcania | 2,955,432 | 3,023,439 | 1,260,938 |
|  | Ngardetta | 1,748,050 | 1,938,258 | 1,010,312 |
| Migration rates | |  |  |  |
|  | me arc/gar | 3.99e-7 | 4.79e-7 | 9.71e-08 |
|  | me gar/arc | 9.07e-8 | 1.11e-7 | 7.27e-08 |

**Table S1:** Results of the demographic analyses conducted with Fastsimcoal2 including or not the Z chromosome in the input file (2D-SFS).

Supplemental Figure S2:

**Patterns of genetic diversity point towards a strong genome-wide differentiation with increased signals on the Z chromosome**

The high sequence coverage along most of the reference genome allowed the estimation of genetic diversity indices for 9,844 abutting genomic windows of 50 kbp (Figure 2). Genetic differentiation (*F*_ST_) between *C. arcania* and *C. gardetta* was high, on average*,* with a mean value of 0.374 when considering the entire genome and 0.357 when excluding the sex chromosome Z (Figure 2). The Z chromosome displayed a strikingly higher level of differentiation than the autosomes, with a mean *F*_ST(Z)_ of 0.687. *F*_ST_ estimates were variable along the chromosomes with a standard deviation of 0.13 and, again, a difference between the autosomes (SD = 0.102) and the Z chromosome, the latter showing higher variability (SD = 0.114). Multiple autosomal genomic regions displayed high values of *F*_ST_ (> 0.6), and the Z chromosome almost uniformly showed *F*_ST_ values superior to 0.6, with a maximum of 0.9 (Figure 2). The absolute genetic divergence *d*_XY_ was negatively correlated with *F*_ST_. This was especially clear on the Z chromosome where the mean *d*_XY_ (0.009) was much lower than on the autosomes (0.018). Genetic diversity estimates (π and *θ*_w_) followed a similar trend as *d*_XY_ with a mean π of 0.014 in *C. arcania* and 0.013 in *C. gardetta* and a mean *θ*_w_ of 0.017 in *C. arcania* and 0.014 in *C. gardetta*. The extremities of the chromosomes displayed increased values of π and *θ*_w_ and lower values of *F*_ST_, suggesting a link between the position of the loci along the chromosomes and their genomic diversity and degree of differentiation between species (fig. 2). Tajima’s *D* was relatively homogeneous along the autosomes, with a mean of -0.65 in *C. arcania* and -0.36 in *C. gardetta*, but very variable on the Z chromosome (mean = -0.57, SD = 0.43 in *C. arcania* and mean = -0.58, SD = 0.50 in *C. gardetta*). A few restricted autosomal regions exhibited pronounced dips in Tajima’s *D* values: for example a region on chromosome 10 or another on chromosome 15 for both *C. gardetta* and *C. arcania.* These regions all showed increased values of *F*_ST_, potentially indicating directional and/or divergent selection. Finally, the level of linkage disequilibrium, quantified by the average *r*^2^ per nucleotide on each genomic window, were twice as high on the Z chromosome and peaking around regions of low Tajima’s *D* or very high *F*_ST_.

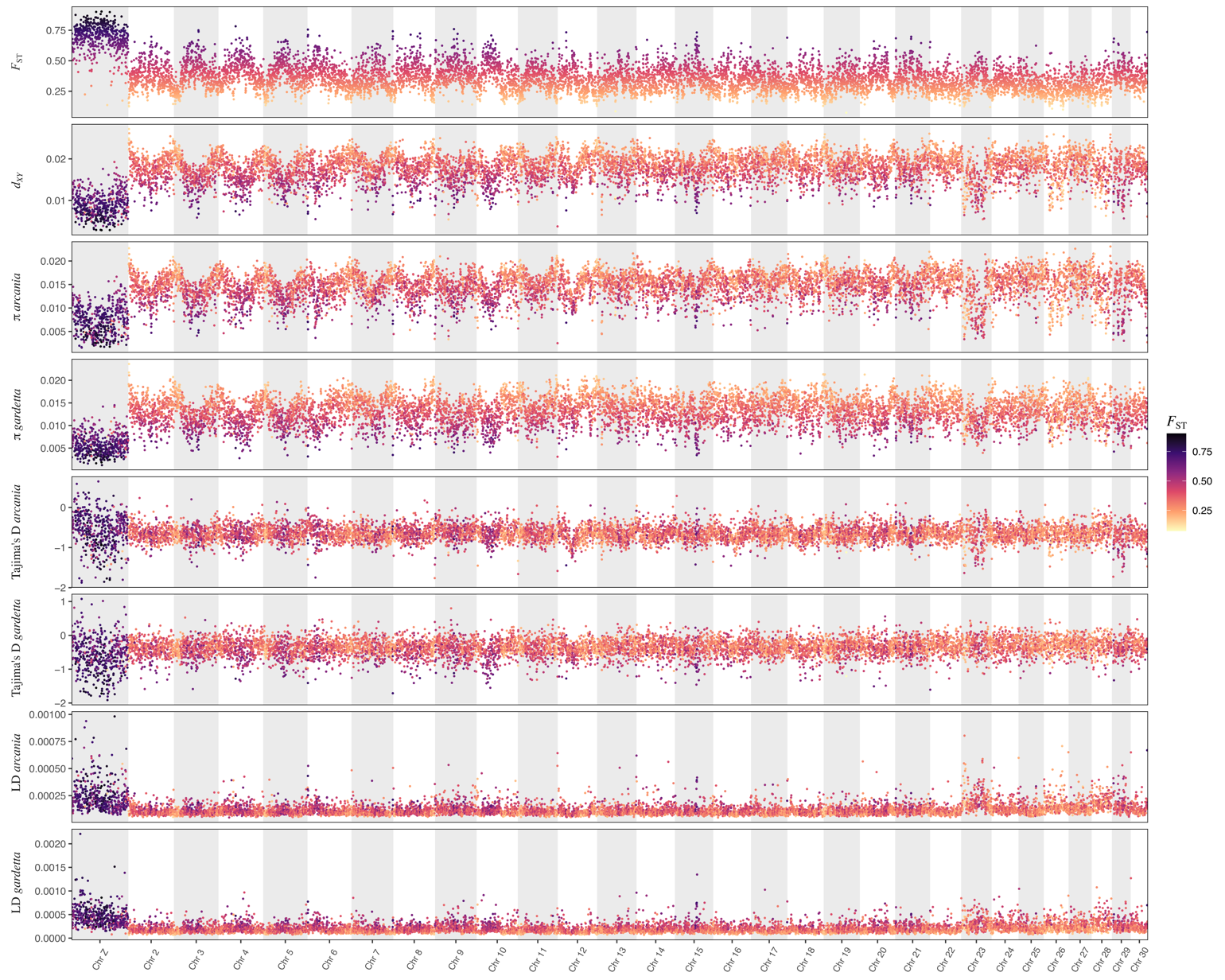

**Fig. S2:** Landscape of genomic diversity and differentiation along *Coenonympha arcania* and *C.* *gardetta* genomes. Species-specific nucleotide diversity (π), Tajima’s *D* and LD are shown together with *F*_ST_ and *d*_XY_ between the two species for all 50kbp abutting genomic windows for which more than 5,000 sites were covered. The color of the dots corresponds to the values of *F*_ST_; they are similar for all subplots and are meant to show correlations between *F*_ST_ and other parameters.

Supplemental Figure S3:

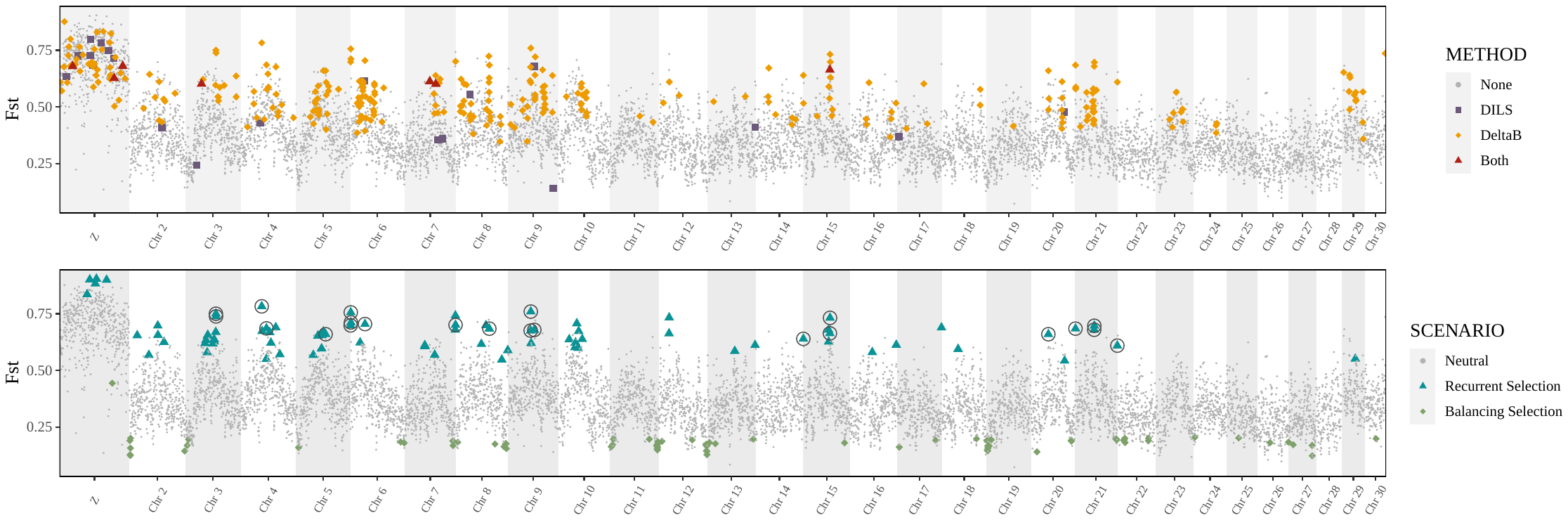

**Fig. S3:** Identification of barriers to gene flow between the genomes of *C. arcania* and *C. gardetta* using a different *m*_e_ value for the Z chromosome (me arc/gar = 3.99e-7, me gar/arc = 9.07e-8) and the autosomes (me arc/gar = 2.32e-04, me gar/arc = 9.26e-05). Genomic windows considered as resistant to gene flow using two process-based procedures: the ∆*B* procedure (∆*B* > 0) and/or DILS (posterior probability of being a barrier > 0.7).

Supplemental Figure S4:

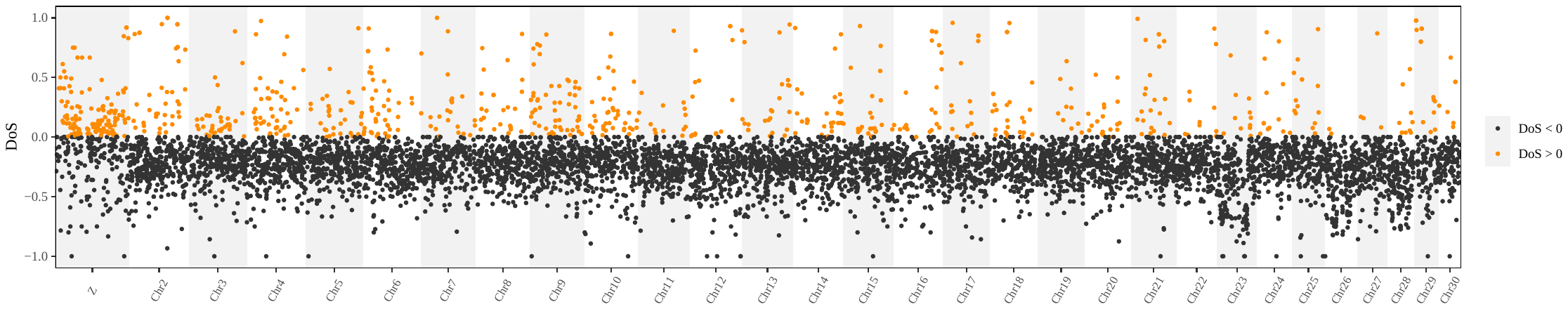

**Fig. S4:** Distribution of Direction of Selection (DoS) along the genome of *Coenonympha*. DoS values were estimated for all 15,239 analyzed genes. DoS values superior to 0, suggesting positive selection, are highlighted in orange.

Supplemental Table S2:

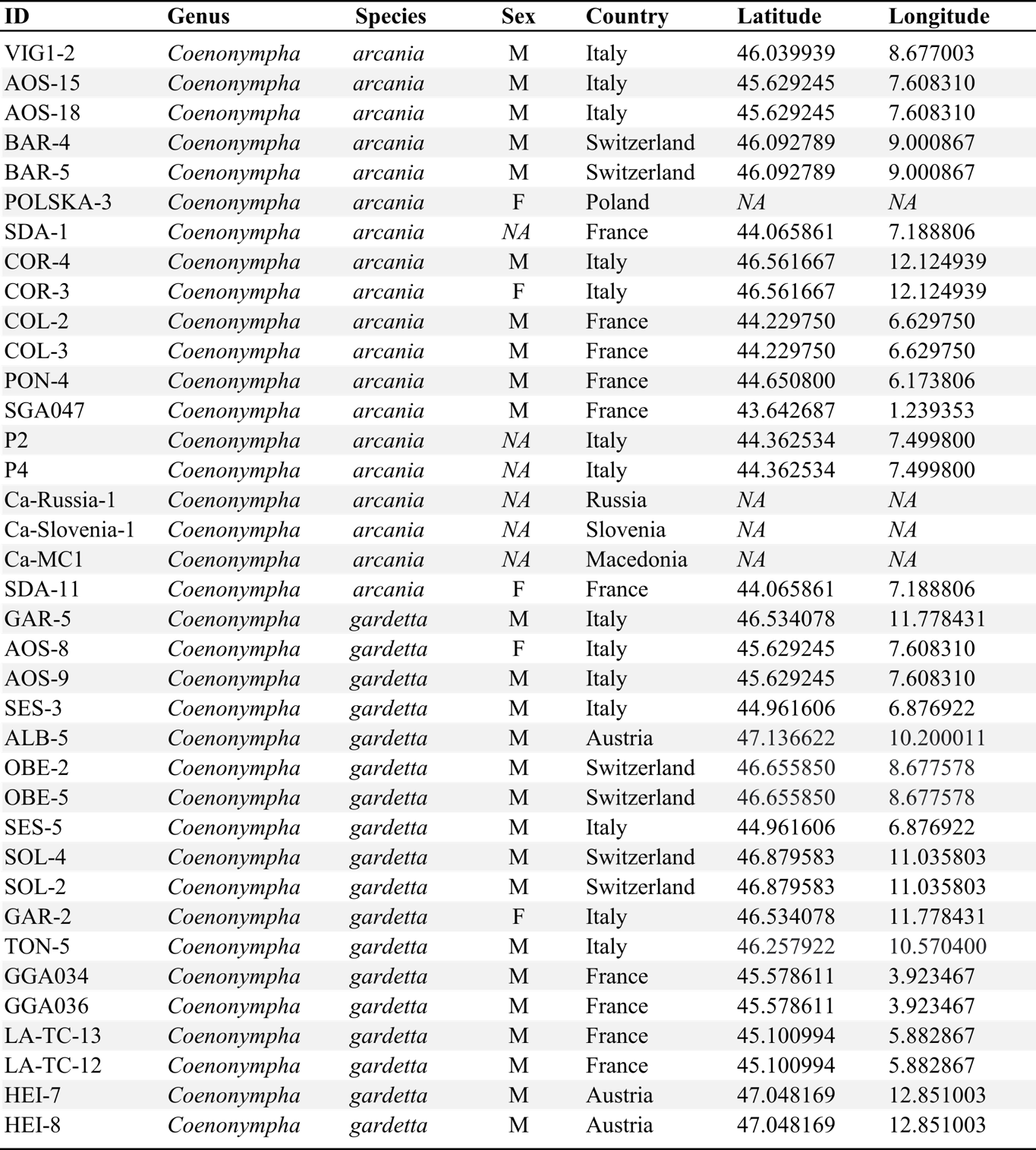

**Table S2:** Characteristics of the 36 sampled *Coenonympha* butterflies.

Supplemental Code S1:

**Genomic data acquisition and processing**

Detailed scripts are also available on Github: https://github.com/Capblancq/Speciation-Coenonympha-butterflies/Speciation-genomics-arcania-gardetta/

*Genotype likelihoods estimation for each population using ANGSD*

Genotype likelihoods were estimated using the SAMtools genotype likelihood model, only using reads having unique best hits (“-uniqueOnly 1”), setting a minimum MapQ score to keep a read to 20 (“-minMapQ 20”), a min nucleotide Q score to consider a site to 20 (“-minQ 20”), a minimum number of 5 individuals with coverage to consider a site (“-minInd 5”), a minimum of 3 and maximum of 60 reads to estimate genotype likelihood for one individual (“-setMinDepthInd 3” and “-setMaxDepthInd 60”), keeping only biallelic sites (“-skipTriallelic 1”), performing the base alignment quality (BAQ: Phred-scaled probability of a read base being misaligned)(Li, 2011) as in SAMtools (“-baq 1”). The command was run as followed:

```
Example for one population:
angsd -b arcania_bam.list \
-ref BST1.fa \
	-anc BST1.fa \
	-out arcania_allsites \
	-nThreads 10 \
	-uniqueOnly 1 -remove_bads 1 -trim 1 -C 50 -baq 1 -minMapQ 20 -minQ 20 \
	-setMinDepthInd 4 -setMaxDepthInd 80 -minInd 5 \
	-skipTriallelic 1 \
	-GL 1 \
	-doMajorMinor 1 -doMaf 1 \
	-doCounts 1
```

We then identified the genomic sites (monomorphic and polymorphic) that were covered for both species as follow:

zcat arcania_allsites.mafs.gz | awk 'BEGIN { OFS = ":" }{ print $1,$2 }' | sed '1d' | sort > arcania_allsites_sites.txt

zcat gardetta_allsites.mafs.gz | awk 'BEGIN { OFS = ":" }{ print $1,$2 }' | sed '1d' | sort > gardetta_allsites_sites.txt

comm -12 arcania_allsites_sites.txt gardetta_allsites_sites.txt > arcania_gardetta_allsites_sites.txt

sed 's/:/\t/' arcania_gardetta_allsites_sites.txt | sort -b -k1,1 > intersect_arcgar_allsites.txt

cut -f1 intersect_arcgar_allsites.txt | uniq | sort > intersect_arcgar_allsites.chr

angsd sites index ${output}/intersect_arcgar_allsites.txt

And re-ran ANGSD with all samples using the “-site” option to produce a global dataset:

```
angsd -b arcania_bam.list \
	-ref BST1.fa \
	-anc BST1.fa \
	-out arcania_allsites \
	-sites intersect_arcgar_allsites.txt \
	-nThreads 10 \
	-uniqueOnly 1 -remove_bads 1 -trim 1 -C 50 -baq 1 -minMapQ 20 -minQ 20 \
	-GL 1 \
	-doMaf 1 -doMajorMinor 1 -doSaf 1 \
#-snp_pval 1e-6 (if only polymorphic sites)
```

*Nucleotide diversity (π) and Tajima’s D*

We used thetaStat, a subprogram of ANGSD, to estimate, for each species various genetic diversity parameters on 50 kbp genomic windows, including nucleotide diversity (π) and Tajima’s D. To do so, we used Site Frequency Spectra (SFS) estimated from the genotype likelihoods dataset (-doSaf 1 option above) and the thetaStat program as follow:

realSFS arcania_allsites.saf.idx -maxIter 50000 -tole 1e-6 -P 10 > arcania.sfs

realSFS saf2theta arcania_allsites.saf.idx -outname arcania -sfs arcania.sfs

thetaStat do_stat arcania.thetas.idx -win 50000 -step 50000 -type 0 -outnames arcania

It is important to note that we used all the covered sites to estimate π and Tajima’s D, including polymorphic and monomorphic sites.

*Pairwise F_ST_ estimation using realSFS*

Pairwise *F*_ST_ were estimated for each genomic window using the program realSFS. To do so, we first produced an optimized Site Frequency Spectrum (SFS) for each population, from the pruned and polymorphic genotype likelihoods dataset:

realSFS arcania_allsites.saf.idx gardetta_allsites.saf.idx -maxIter 50000 -tole 1e-6 -P 10 > arcania.gardetta.sfs

Then, we used the optimized SFS of each population to produce 2D SFS for each pair of population and estimated *F*_ST_ from these 2D SFS:

```
realSFS fst index ${input}/arcania_allsites.saf.idx gardetta_allsites.saf.idx \
	-sfs arcania.gardetta.sfs \
	-fstout arcania-gardetta_Fst
```

And obtained the estimate along sliding windows:

realSFS fst stats2 arcania-gardetta_Fst.fst.idx -win 50000 -step 50000 -type 0 > Fst_sliding_arcania-gardetta.txt

Supplemental Table S3:

**Testing the goodness-of-fit of the ABC procedure in DILS**

Goodness-of-fit tests are performed by simulating under the best model each population genetic statistic. Genetic data were simulated under the best supported model, using the estimated parameters. These simulations then empirically produce the statistical distributions summarized under the inferred model. We then examine whether the value observed from the *Coenonympha* dataset is correctly captured by the inferred model for each of the summary statistics.

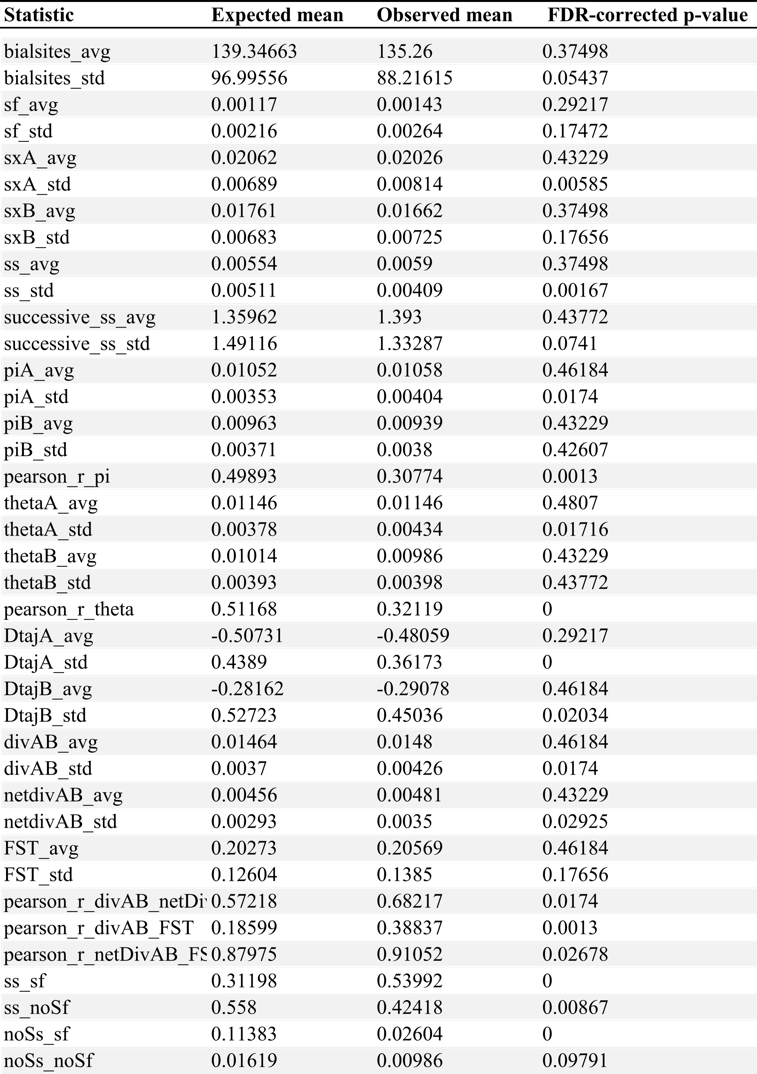

**Table S3:** Goodness-of-fit for each summary statistic with the expected mean, observed mean, and the p-value corrected for multiple testing. The expected values here are based on the optimized posterior simulations and p-values > 0.01 show that simulations made with the optimized parameter values can accurately reproduce the statistic value obtained with the observed data.

Supplemental Figure S4:

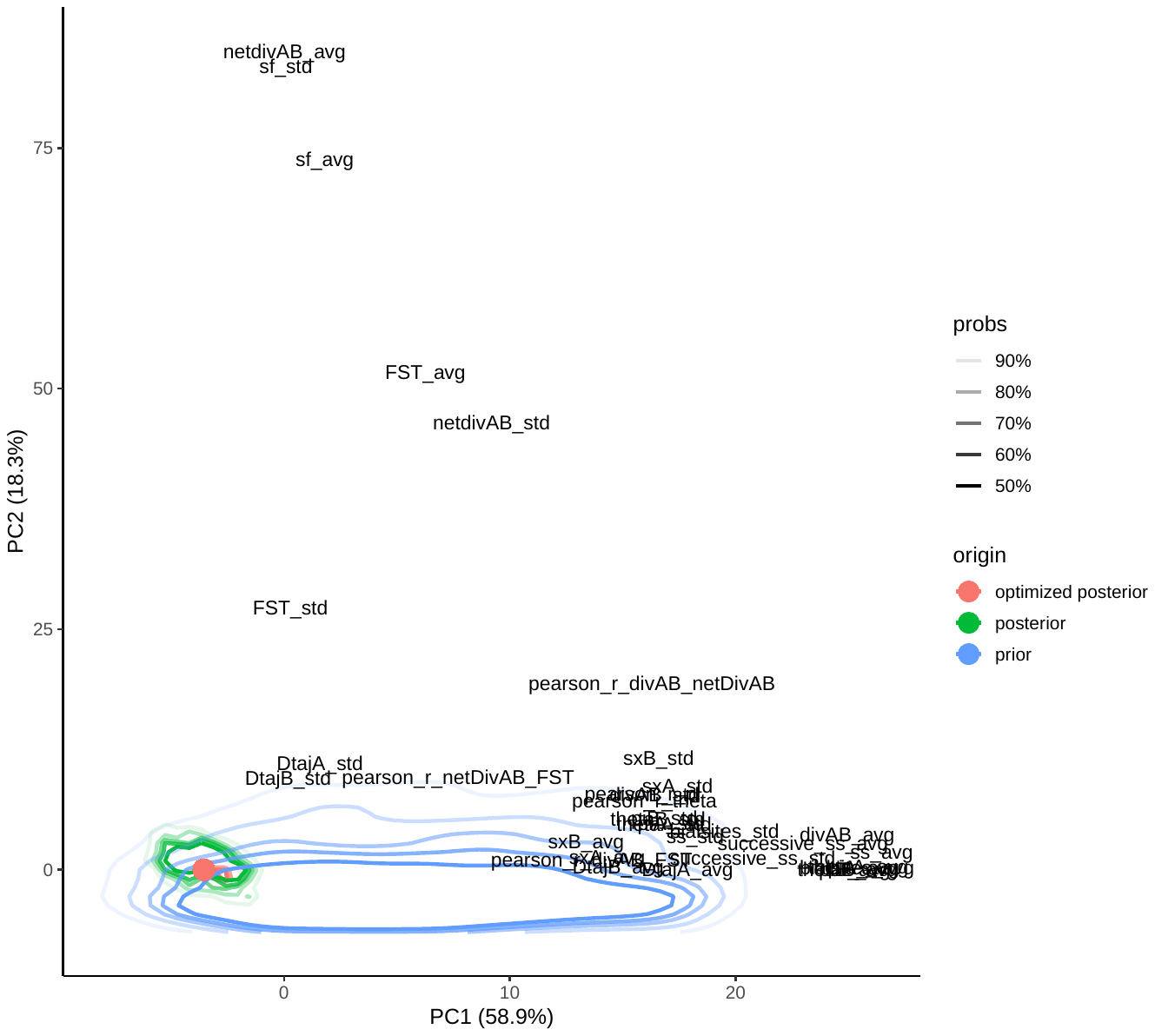

**Fig. S4:** Principal component analysis showing the match between the observed dataset (red dot) and simulations produced using the optimized posterior parameter values. The distributions of 2,000 prior simulations are shown in blue, 2,000 posterior simulations in green and 2,000 optimized simulations in red, together with the influence of each statistic in the variation.

Supplemental Code S2:

**Identifying barrier loci using DILS and gIMble**

Detailed scripts are also available on Github: https://github.com/Capblancq/Speciation-Coenonympha-butterflies/Speciation-genomics-arcania-gardetta/

*Generating input file for DILS*

Run for each genomic window considered, the following code allows to produce the alignment required as input file for DILS.

```
# Extract chrom and pos of the target window
CHROM=$(chromosome considered)
POS=$(midPos of the window)
POS1=$((${POS}-25000))
POS2=$((${POS}+25000))

# Create a file with information requires for the header of DILS’s input file
touch names.tmp
for i in $(cat bam.list)
do
	name=${i/.bam/}
	echo `basename $name` >> names.tmp
done
paste -d "|" species.txt names.tmp > header.tmp
cat header.tmp | tr '\n' ' ' | sed -e 's/ /\t/g' > header.txt
rm *.tmp

# Produce a geno file with genotypes for all individuals and positions (including monomorphic) for the genomic window
angsd -b all_bam.list \
    -ref BST1.fa \
    -anc BST1.fa \
    -out window \
    -r ${CHROM}:${POS1}-${POS2} \
    -nThreads 10 \
    -uniqueOnly 1 -remove_bads 1 -trim 1 -C 50 -baq 1 -minMapQ 20 -minQ 20 \
    -GL 1 \
    -doMaf 1 -doMajorMinor 1 \
    -doPost 1 -postCutoff 0.7 -doGeno 4 -doCounts 1 -geno_minDepth 3 -geno_maxDepth 200

# Check the number of sites genotypes
nb_loci=$(zcat window.geno.gz | wc -l)

if (("${nb_loci}" < "5000"))
then

	echo "Not enough sites on window"

else
# Format the file
zcat window.geno.gz | cut -f3- > window.tmp
cat header.txt <(echo) window.tmp > geno_window.tmp

# Dissociate the two alleles and produce the sequence alignment
cat geno_window.tmp | gawk '{gsub(/\t$/,"");}1' | gawk -F"\t" 'FNR==1{for (i=1;i<=NF;i++) {header[i] = $i;}nbInd=NF;next;}{for (i=1;i<=nbInd;i++) {seq[i,1] = seq[i,1] "" substr($i,1,1);seq[i,2] = seq[i,2] "" substr($i,2,1);}}END{for (i=1;i<=nbInd;i++) {for (j=1;j<=2;j++) print ">"header[i]"|allele"j"\n" seq[i,j]}}' > window.dils

	# Complete the header for each allele
  	sed -i "s/>/>window_${CHROM}:${POS}|/g" window.dils

fi

# Concatenate all the files together (has to be done once all the window files are done!)
cat *.dils > input.dils.fasta
```

*Running DILS*

DILS was run using snakemake and a config file containing multiple filtering threshold options such as a maximum percentage of N tolerated (max_N_tolerated=0.75), a minimum number of sequences to consider a locus (nMin=6) or a specific mutation rate (mu=2.9*10-9).

```
# Config file
mail_address:
infile: input.dils.fasta
region: noncoding
nspecies: 2
nameA: arcania
nameB: gardetta
nameOutgroup: NA
lightMode: TRUE
useSFS: 1
config_yaml: myConfigFileTwoPops.yaml
timeStamp: DILS_output
population_growth: variable
modeBarrier: bimodal
max_N_tolerated: 0.75
Lmin: 100
nMin: 6
mu: 0.0000000029
rho_over_theta: 0.5
N_min: 0
N_max: 5000000
Tsplit_min: 100000
Tsplit_max: 3000000
M_min: 1
M_max: 40

# Runs DILS
snakemake --snakefile Snakefile_2pop \
	-p \
	-j 140 \
	--configfile myConfigFileTwoPops_noZ.yaml \
	--cluster-config cluster_2pop.json \
	--cluster "sbatch --nodes={cluster.node} --ntasks={cluster.n} --cpus-per-task={cluster.cpusPerTask} --time={cluster.time} --mem-per-cpu=30G" \
	--latency-wait 60 \
	--rerun-incomplete
```

*Generating input file for our gIMble-like procedure*

Run for each genomic window considered, the following code allows to produce the 2D-SFS required as input file for the gIMble-like analysis we conducted using Fastsimcoal2.

```
# Extract chrom and pos of the target window
CHROM=$(chromosome considered)
POS=$(midPos of the window)
POS1=$((${POS}-25000))
POS2=$((${POS}+25000))

# Estimates 2D-SFS from saf files
realSFS arcania_allsites.saf.idx gardetta_allsites.saf.idx -r ${CHROM}:${POS1}-${POS2} > window.sfs

# Creates 2D-SFS with Fastsimcoal format arcania (pop0) / gardetta (pop1)
R –vanilla << EOF

sfs <- scan(paste(“window.sfs”, sep=""), quiet=T)
N1 <- 19
N2 <- 16
tab <- matrix(data = sfs, ncol = (N1*2+1), nrow = (N2*2+1))
colnames(tab) <- paste(rep("d0", (N1*2+1)), seq(0,(N1*2)), sep = "_")
row.names(tab) <- paste(rep("d1", (N2*2+1)), seq(0,(N2*2)), sep = "_")

# Function to transform dadi like SFS into 2D-SFS for Fastsimcoal2
derived2maf=function(derived_sfs2d){
		n1=nrow(derived_sfs2d)
	 	n2=ncol(derived_sfs2d)
	 	maf_2dsfs=matrix(0,nrow=n1,ncol=n2)
	 	colnames(maf_2dsfs)=colnames(derived_sfs2d)
	 	rownames(maf_2dsfs)=rownames(derived_sfs2d)
	 	threshold_freq=0.5*(n1+n2-2)
	 	for(i in 0:(n1-1)){
	   	for(j in 0:(n2-1)){
	     	if(i+j < threshold_freq){
	       	maf_2dsfs[i+1,j+1]=maf_2dsfs[i+1,j+1]+derived_sfs2d[i+1,j+1]
	     	}
	     	else if(i+j == threshold_freq){
	       maf_2dsfs[i+1,j+1]=maf_2dsfs[i+1,j+1]+(0.5*derived_sfs2d[i+1,j+1]+0.5*derived_sfs2d[n1-i,n2-j])
	     	}
	     	else{
	       	maf_2dsfs[n1-i,n2-j]=maf_2dsfs[n1-i,n2-j]+derived_sfs2d[i+1,j+1]
	     	}
	   	}
	 	}
	 	maf_2dsfs
}

# Convert SFS
tab2 <- derived2maf(tab)

# Write output with the right name for Fastsimcoal2
write.table(tab2, “window_jointMAFpop1_0.obs”, row.names = T, col.names = T, sep = "\t", quote = F)

EOF

# Add tabulation first line
sed -i 's/d0_0/\td0_0/g' window_jointMAFpop1_0.obs

# Add header
echo "1 observations" > header.txt
cat header.txt window_jointMAFpop1_0.obs > window_jointMAFpop1_0.tmp
mv window_jointMAFpop1_0.tmp window_jointMAFpop1_0.obs
```

*Running the gIMble-like procedure*

Fastsimcoal2 was run on the 2D-SFS using two different models of divergence, either with or without gene flow between species (see the two template files below). The likelihood of the two models optimized with the observed window 2D-SFS were then compared to decide on the presence of gene flow.

```
# Creates directory for the scenario
mkdir window

# Copies required files
cp SC_windows_m0.tpl SC_windows_m0.est window/
cp window_jointMAFpop1_0.obs window/SC_windows_m0_jointMAFpop1_0.obs
cp SC_windows_mfixed.tpl SC_windows_mfixed.est window/
cp window_jointMAFpop1_0.obs window/SC_windows_mfixed_jointMAFpop1_0.obs

# Enters the run directory
cd window/

# Run fastsimcaol with varying me
fsc2705 -t SC_windows_m0.tpl -n1000000 -m -e SC_windows_m0.est -M -L40 -q -c10

# Run Fastsimcoal with fixed me
fsc2705 -t SC_windows_mfixed.tpl -n1000000 -m -e SC_windows_mfixed.est -M -L40 -q -c10

# Estimate delta B following Laetsch et al. 2022
lnCL1=`cat window/SC_windows_m0/SC_windows_m0.bestlhoods | awk 'NR==2 {print $3}'`
lnCL2=`cat window/SC_windows_mfixed/SC_windows_mfixed.bestlhoods | awk 'NR==2 {print $3}'`
DeltaB=$(bc <<< "${lnCL1} - ${lnCL2}")

# Appends results to the results file
if test -f results_SC_m0_windows.txt
then
	echo "${CHROM} ${POS}" `cat window/SC_windows_m0/SC_windows_m0.bestlhoods | awk 'NR==2'` ${DeltaB} >> results_SC_m0_windows.txt
else
	echo "Chrom midPos" `cat window/SC_windows_m0/SC_windows_m0.bestlhoods | awk 'NR==1'` "DeltaB" > results_SC_m0_windows.txt
	echo ${CHROM} ${POS} `cat window/SC_windows_m0/SC_windows_m0.bestlhoods | awk 'NR==2'` ${DeltaB} >> results_SC_m0_windows.txt
fi

##### FPR estimation #####

# Retreives .par file of the best run of the best SC scenario for the divergence of arcania and gardetta
cp ./SC_windows_mfixed/SC_windows_mfixed_maxL.par ./
sed -i 's/^1 0$/200 0/g' SC_windows_mfixed_maxL.par
sed -i 's/^FREQ 1/DNA 100/g' SC_windows_mfixed_maxL.par

# Then generate 100 SFS
~/TOOLS/fsc2705 -i SC_windows_mfixed_maxL.par -n100 -j -d -s0 -x -I -q -c10 -m

# Prepares file to store the results
echo "Run" "DeltaB" > ./results_FPR_${CHROM}_${POS}.txt

# Running the search for barrier loci with each SFS
for i in {1..100}
do
	cd ./SC_windows_mfixed_maxL/SC_windows_mfixed_maxL_${i}/

	# Run fastsimcaol with me 0
	cp ${OUTPUT}/SC_windows_m0.tpl ./SC_windows_m0_maxL.tpl
	cp ${OUTPUT}/SC_windows_m0.est ./SC_windows_m0_maxL.est
	cp SC_windows_mfixed_maxL_jointMAFpop1_0.obs SC_windows_m0_maxL_jointMAFpop1_0.obs
	~/TOOLS/fsc2705 -t SC_windows_m0_maxL.tpl -n10000 -m -e SC_windows_m0_maxL.est -M 0.01 -L40 -q -c10

	# Run fastsimcaol with fixed me
	cp ${OUTPUT}/SC_windows_mfixed.tpl ./SC_windows_mfixed_maxL.tpl
	cp ${OUTPUT}/SC_windows_mfixed.est ./SC_windows_mfixed_maxL.est
	~/TOOLS/fsc2705 -t SC_windows_mfixed_maxL.tpl -n10000 -m -e SC_windows_mfixed_maxL.est -M 0.01 -L40 -q -c10

	# Estimate delta B following Laetsch et al. 2022
	lnCL1=`cat ./SC_windows_m0_maxL/SC_windows_m0_maxL.bestlhoods | awk 'NR==2 {print $3}'`
	lnCL2=`cat ./SC_windows_mfixed_maxL/SC_windows_mfixed_maxL.bestlhoods | awk 'NR==2 {print $3}'`
	DeltaB=$(bc <<< "${lnCL1} - ${lnCL2}")

	# Appends results to the results file
	echo ${i} ${DeltaB} >> ../../results_FPR_${CHROM}_${POS}.txt

	# Go back to window directory
	cd ../../

done

# Count number of dletaB values above 0 to estimate FPR per window
FPR=`awk 'FNR==1 {count = 0; next} $2 > 0 {count++} END {print (count+0.0)/(NR-1);}' results_FPR_${CHROM}_${POS}.txt`

# Append value to the result file
if test -f ${OUTPUT}/results_FPR_windows.txt
then
	echo ${CHROM} ${POS} ${FPR} >> ${OUTPUT}/results_FPR_windows.txt
else
	echo "CHROM" "POS" "FPR" > ${OUTPUT}/results_FPR_windows.txt
	echo ${CHROM} ${POS} ${FPR} >> ${OUTPUT}/results_FPR_windows.txt
fi


# .tpl file for Fastsimcoal2 with no geneflow
//Number of population samples (demes)
2
//Population effective sizes (number of genes)
N1
N2
//Sample sizes
38
32
//Growth rates : negative growth implies population expansion
0
0
//Number of migration matrices : 0 implies no migration between demes
0
//historical event: time, source, sink, migrants, new size, new growth rate, migr. matrix
1 historical event
1793459 1 0 1 1 0 0
//Number of independent loci [chromosome]
1 0
//Per chromosome: Number of linkage blocks
1
//per Block: data type, num loci, rec. rate and mut rate + optional parameters
FREQ 1 0 2.9e-9 OUTEXP

# .tpl file for Fastsimcoal2 with geneflow
//Number of population samples (demes)
2
//Population effective sizes (number of genes)
N1
N2
//Sample sizes
38
32
//Growth rates  : negative growth implies population expansion
0
0
//Number of migration matrices : 0 implies no migration between demes
2
//Migration matrix 0
0 9.07553e-8
3.99313e-7 0
//Migration matrix 1
0 0
0 0
//historical event: time, source, sink, migrants, new size, new growth rate, migr. matrix
2 historical events
360240 0 0 0 1 0 1
1793459 1 0 1 1 0 1
//Number of independent loci [chromosome]
1 0
//Per chromosome: Number of linkage blocks
1
//per Block: data type, num loci, rec. rate and mut rate + optional parameters
FREQ 1 0 2.9e-9 OUTEXP

# .est file
// Priors and rules file
// *********************
[PARAMETERS]
//#isInt? #name   #dist.#min  #max
//all Ns are in number of haploid individuals
1	N1	unif	1000000	4000000	output
1	N2	unif	500000	3000000	output
[RULES]
[COMPLEX PARAMETERS]
```
